# Sequential compression of gene expression across dimensionalities and methods reveals no single best method or dimensionality

**DOI:** 10.1101/573782

**Authors:** Gregory P. Way, Michael Zietz, Vincent Rubinetti, Daniel S. Himmelstein, Casey S. Greene

## Abstract

**Background:** Unsupervised compression algorithms applied to gene expression data extract latent, or hidden, signals representing technical and biological sources of variation. However, these algorithms require a user to select a biologically-appropriate latent dimensionality. In practice, most researchers select a single algorithm and latent dimensionality. We sought to determine the extent by which using multiple dimensionalities across ensemble compression models improves biological representations.

**Results:** We compressed gene expression data from three large datasets consisting of adult normal tissue, adult cancer tissue, and pediatric cancer tissue. We compressed these data into many latent dimensionalities ranging from 2 to 200. We observed various tradeoffs across latent dimensionalities and compression models. For example, we observed high model stability between principal components analysis (PCA), independent components analysis (ICA), and non-negative matrix factorization (NMF). We identified more unique biological signatures in ensembles of denoising autoencoder (DAE) and variational autoencoder (VAE) models in intermediate latent dimensionalities. However, we captured the most pathway-associated features using all compressed features across algorithms and dimensionalities. Optimized at different latent dimensionalities, compression models detect generalizable gene expression signatures representing sex, neuroblastoma MYCN amplification, and cell types. In two supervised machine learning tasks, compressed features optimized predictions at different latent dimensionalities.

**Conclusions:** There is no single best latent dimensionality or compression algorithm for analyzing gene expression data. Instead, using feature ensembles from different compression models across latent space dimensionalities optimizes biological representations.

## Background

Dimensionality reduction algorithms compress input data into feature representations that capture sources of variation. Applied to gene expression data, compression algorithms can identify latent biological representations and technical processes. These biological representations reveal important information about the samples and can help to generate hypotheses that are difficult or impossible to observe in the original genomic space. For example, applying PCA to a large cancer transcriptomic compendium determined the influence of copy number alterations on gene expression measurements [1]. Applying ICA to transcriptome data aggregated gene modules representing core pathways and hidden transcriptional programs [2, 3]. Training NMF models using bulk gene expression data estimated cell type proportion [4, 5]. DAEs have revealed latent signals characterizing oxygen exposure and transcription factor targets [6, 7], and VAEs have identified biologically relevant latent features that discriminate cancer subtypes and drug response [8, 9]. Additional latent variable approaches have been used to detect and remove technical artifacts, including batch effects [10, 11]. Here, we focus on using compression to identify biological representations by analyzing processed data with batch effect already mitigated. Nevertheless, a major challenge to all compression applications is the fundamental requirement that a researcher must determine the number of latent dimensions (*k*) to compress input data.

Instead, it is possible that different biological signatures are best captured at different latent space dimensionalities. To test this, we trained and evaluated various compression models across a wide range of latent space dimensionalities, from *k* = 2 to *k* = 200. We train PCA, ICA, NMF, DAE, and VAE models using RNAseq gene expression data. We selected these methods because they are either widely established in practice (PCA, NMF) or use neural networks that are rapidly growing in popularity (DAE, VAE). We included ICA to compare against PCA: it has been used in practice [7] and also allows us to compare multiple rotations of the low-dimensional space. We applied these methods to three different datasets. We applied these methods to three different datasets: The Cancer Genome Atlas (TCGA) PanCanAtlas [12], the Genome Tissue Expression Consortium Project (GTEx) [13], and the Therapeutically Applicable Research To Generate Effective Treatments (TARGET) Project [14].

We demonstrate various model tradeoffs in reconstruction cost, stability, and gene set coverage in training and testing sets across datasets, algorithms, and latent dimensionalities. We observe that several distinct gene expression signatures are optimized in various models spanning low, intermediate, and high latent dimensionalities. Our primary finding is that there is no single algorithm or dimensionality that is best for all purposes: Instead, using various latent dimensionalities and algorithms optimizes biological representations. Those who plan to apply these algorithms may consider training multiple models over multiple dimensionalities to avoid missing important biological features.

We name this sequential compression approach “BioBombe” after the large mechanical device developed by Alan Turing and other cryptologists in World War II to decode encrypted messages sent by Enigma machines. Using the BioBombe approach, we compress gene expression input data with multiple latent dimensionalities to decipher and enhance biological representations embedded within compressed gene expression features.

## Results

### BioBombe implementation

We compressed processed RNAseq data from TCGA, GTEx, and TARGET using PCA, ICA, NMF, DAE, and VAE across 28 different latent dimensionalities (*k*) ranging from *k* = 2 to *k* = 200. We split each dataset into 90% training and 10% test sets balanced by cancer type or tissue type and trained models using only the training data. We used real and permuted data and initialized each model five times per latent dimensionality resulting in a total of 4,200 different compression models (**Additional File 1: Figure S1**). We evaluated hyperparameters for DAE and VAE models across dimensionalities and trained models using optimized parameter settings (**Additional File 2**; **Additional File 1: Figure S2**). See **Fig. 1** for an outline of our approach. We provide full BioBombe results for all compression models across datasets for both real [15–17] and permuted data [18–20] in both training and test sets as publicly available resources (see https://greenelab.github.io/BioBombe/).

**Figure 1:**
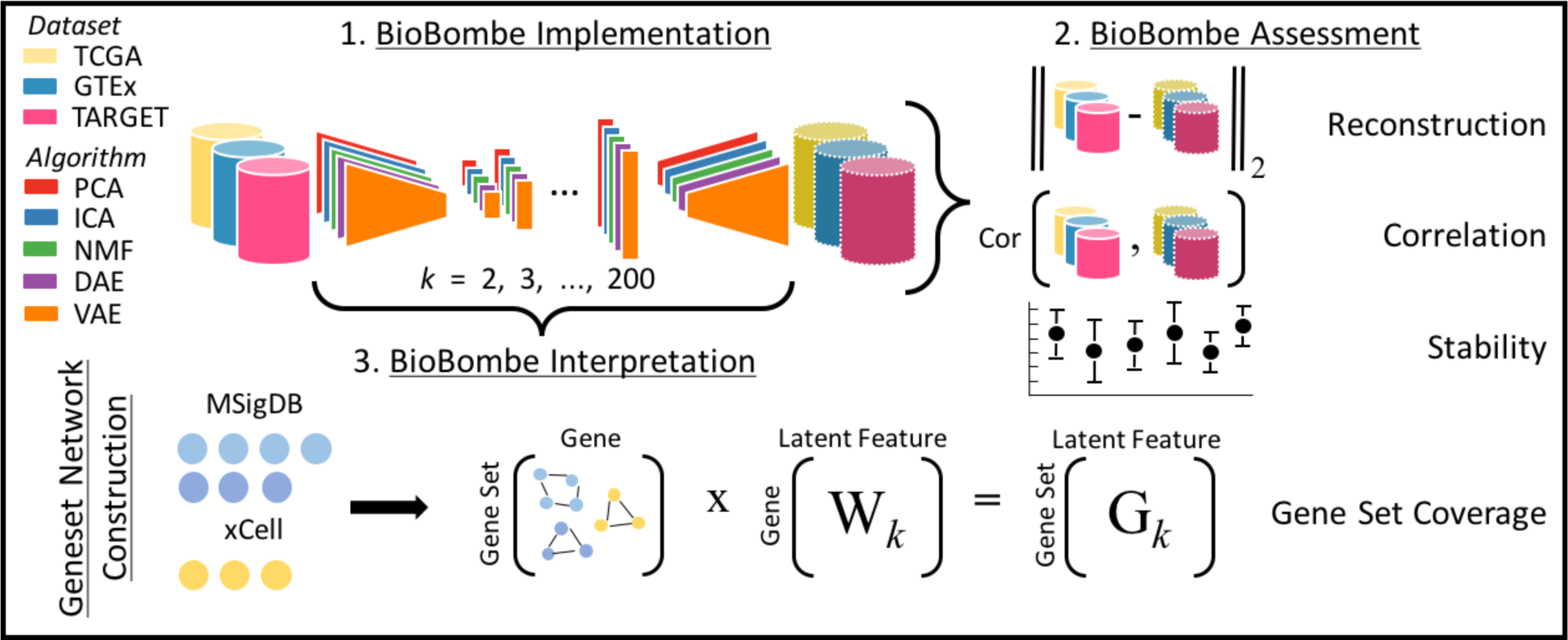
Overview of the BioBombe approach. We implemented BioBombe on three datasets using five different algorithms. We sequentially compressed input data into various latent dimensionalities. We calculated various metrics that describe different benefits and trade-offs of the algorithms. Lastly, we implemented a network projection approach to interpret the compressed latent features. We used MSigDB collections and xCell gene sets to interpret compressed features.

### Assessing compression algorithm reconstruction

Reconstruction cost, a measurement of the difference between the input and output matrices, is often used to describe the ability of compression models to capture fundamental processes in latent space features that recapitulate the original input data. We tracked the reconstruction cost for the training and testing data partitions for all datasets, algorithms, latent dimensionalities, and random initializations. As expected, we observed lower reconstruction costs in models trained with real data and with higher latent dimensionalities (**Additional File 1: Figure S3**). Because PCA and ICA are rotations of one another, we used their identical scores as a positive control. All the compression algorithms had similar reconstruction costs, with the highest variability existing in low latent dimensionalities (**Additional File 1: Figure S3**).

### Evaluating model stability and similarity within and across latent dimensionalities

We applied singular vector canonical correlation analysis (SVCCA) to algorithm weight matrices to assess model stability within algorithm initializations, and to determine model similarity between algorithms [21]. Briefly, SVCCA calculates similarity between two compression algorithm weight matrices by learning appropriate linear transformations and iteratively matching the highest correlating features. Training with TCGA data, we observed highly stable models within algorithms and within all latent dimensionalities for PCA, ICA, NMF (along the matrix diagonal in **Fig 2a**). VAE models were also largely stable, with some decay in higher latent dimensionalities. However, DAE models were unstable, particularly at low latent dimensionalities (**Fig 2a**). We also compared similarity across algorithms. Because PCA and ICA are rotations of one another, we used the high stability as a positive control for SVCCA estimates. NMF was also highly similar to PCA and ICA, particularly at low latent dimensionalities (**Fig. 2a**). VAE models were more similar to PCA, ICA, and NMF than DAE models, particularly at low latent dimensionalities, and the instability patterns within DAE models also led to large differences across algorithms (**Fig. 2a**). We observed similar patterns in GTEx and TARGET data, despite TARGET containing only about 700 samples (**Additional File 1: Figure S4**).

**Figure 2:**
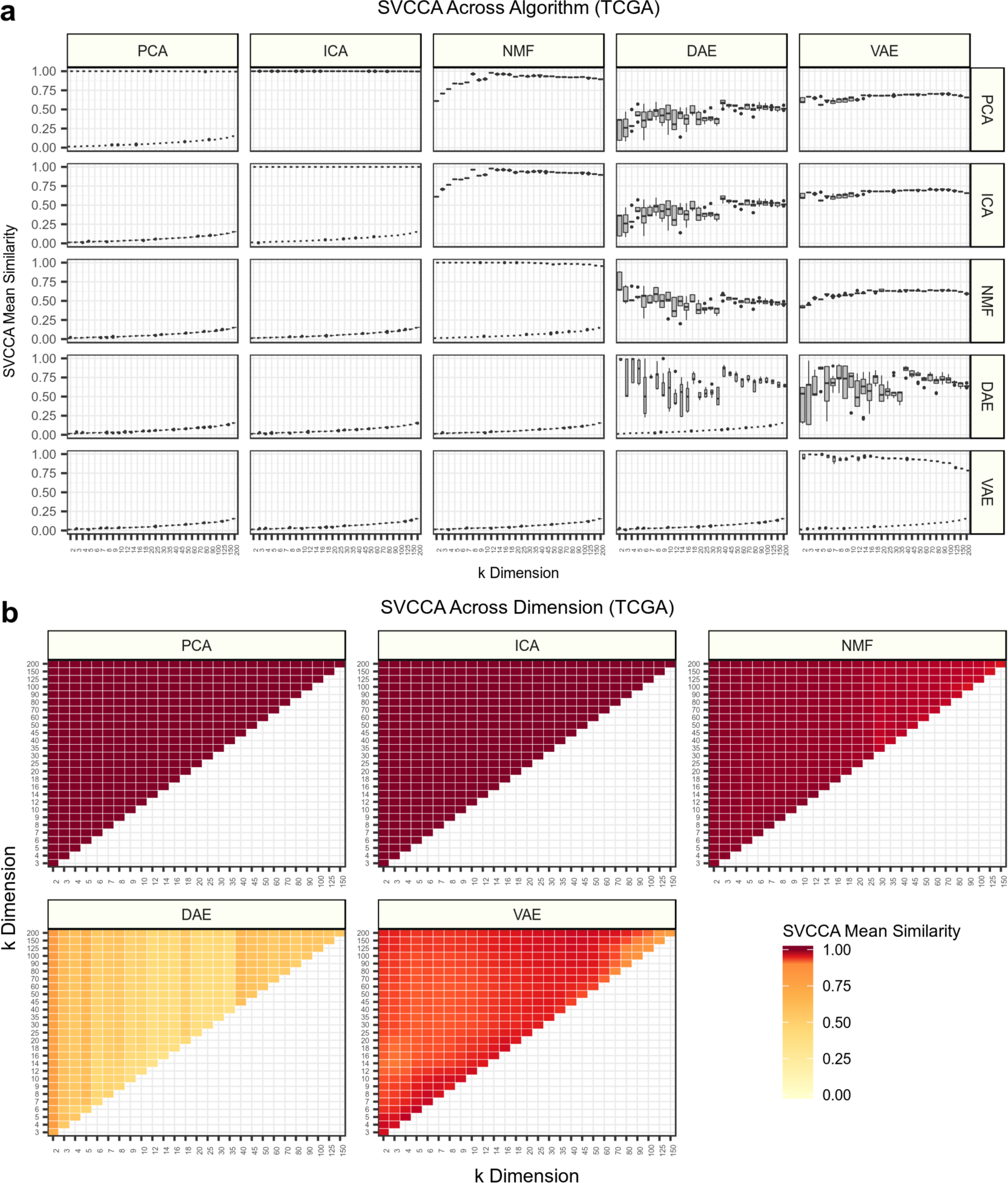
Assessing algorithm and dimensionality stability with singular vector canonical correlation analysis (SVCCA). **(a)** SVCCA applied to the weight matrices learned by each compression algorithm in gene expression data from The Cancer Genome Atlas (TCGA). The mean of all canonical correlations comparing independent iterations is shown. The distribution of mean similarity represents a comparison of all pairwise iterations within and across algorithms. The upper triangle represents SVCCA applied to real gene expression data, while the lower triangle represents permuted expression data. Both real and permuted data are plotted along the diagonal. **(b)** Mean correlations of all iterations within algorithms but across *k* dimensionalities. SVCCA will identify min(i, j) canonical vectors for latent dimensionalities *k_i_* and *k_j_*. The mean of all pairwise correlations is shown for all combinations of *k* dimensionalities.

We also used SVCCA to compare the similarity of weight matrices across latent dimensionalities. Both PCA and ICA found highly similar solutions (**Fig. 2b**). This is expected since the solutions are deterministic and are arranged with decreasing amounts of variance. NMF also identified highly similar solutions in low dimensionalities, but solutions were less similar in higher dimensionalities. DAE solutions were the least similar, with intermediate dimensionalities showing the lowest mean similarity. VAE models displayed relatively high model similarity, but there were regions of modest model stability in intermediate and high dimensionalities (**Fig. 2b**). We observed similar patterns in GTEx and TARGET data (**Additional File 1: Figure S5**).

### BioBombe features capture various gene expression representations across different latent dimensionalities

We tested the ability of features derived using the BioBombe approach to isolate various biological representations. First, we tested the ability to differentiate sample sex; which has been previously observed to be captured in latent space features [8,22,23]. We performed a two-tailed t-test comparing male and female samples in the GTEx test set across all initializations, algorithms, and latent dimensionalities. We optimally identified this phenotype in higher latent dimensionalities, particularly in NMF and VAE models (**Fig. 3a**). The top feature separating GTEx males and females was NMF feature 111 in *k* = 200 (*t* = 44.5, *p* = 7.3 x 10^-176^) (**Fig 3b**). We examined the genes that contribute with high weight to this feature and found only three genes had substantial influence. These three genes all had high positive weights and were encoded on the Y chromosome. We performed the same approach using BioBombe features to identify sex features in TCGA test data (**Fig. 3c**). The top latent dimensionality identified was not consistent across algorithms. The top feature distinguishing TCGA males and females was ICA feature 53 in the *k* = 90 model (*t* = 4.9, *p* = 2.0 x 10^-6^) (**Fig. 3d**). The separation was not as strong using the more complex TCGA data, but the top 10 gene weights were all encoded on the X chromosome.

**Figure 3:**
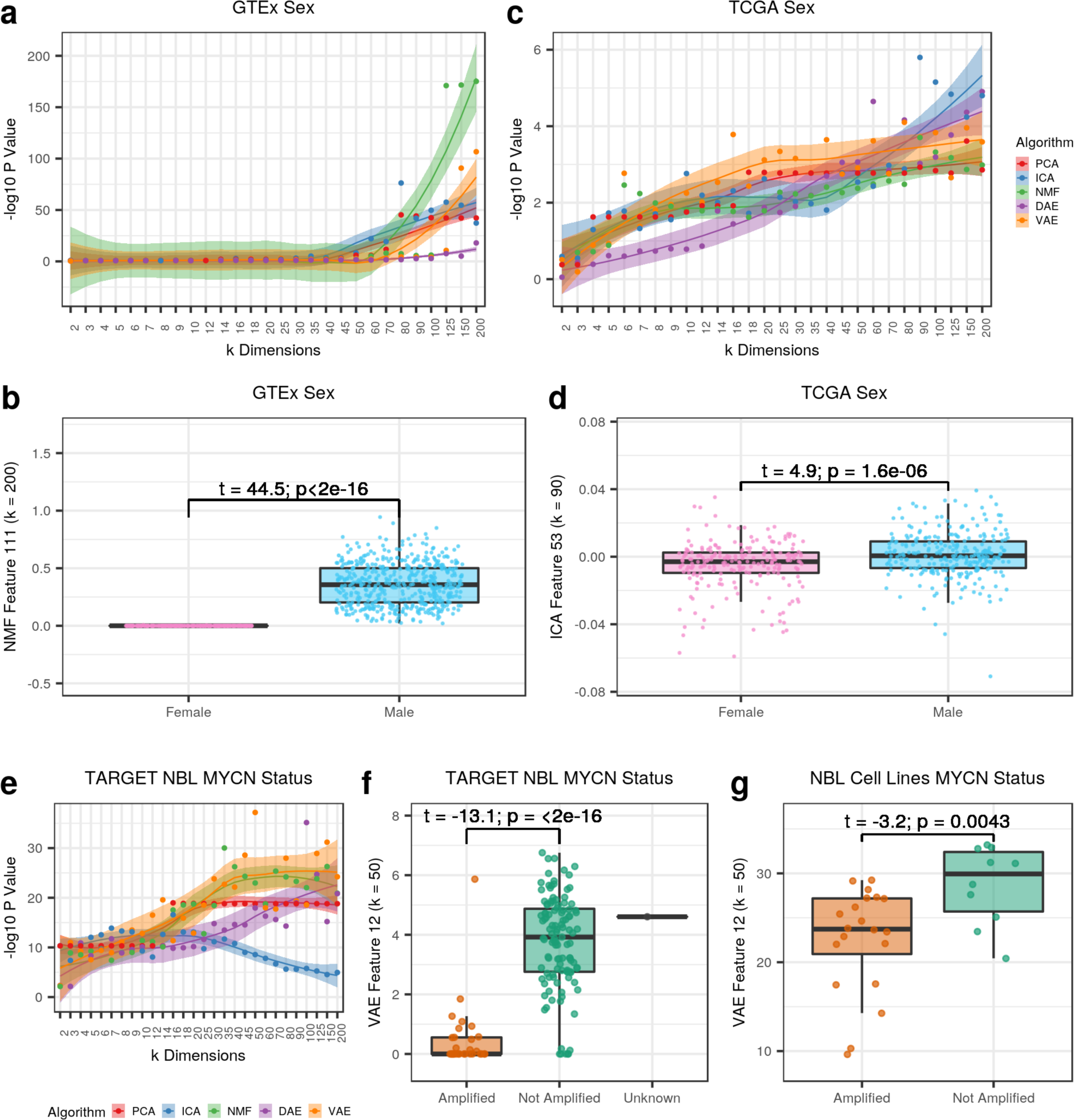
Using BioBombe as a signature discovery tool. Detecting GTEx sample sex across **(a)** various latent dimensionalities and algorithms, and **(b)** the latent feature with the highest enrichment. Detecting TCGA patient sex across **(c)** various latent dimensionalities, and **(d)** the latent feature with the highest enrichment. Detecting TARGET MYCN amplification in neuroblastoma (NBL) tumors **(e)** across various latent dimensionalities, and **(f)** the latent feature with the highest enrichment. **(g)** Applying the MYCN signature to an external dataset of NBL cell lines implicates MYCN amplified cell lines.

We also tested the ability of features derived from the BioBombe approach to distinguish MYCN amplification in neuroblastoma (NBL) tumors. MYCN amplification is a biomarker associated with poor prognosis in NBL patients [24]. We performed a two-tailed t-test comparing MYCN amplified vs. MYCN not amplified NBL tumors for each of the latent features derived from the full set of TARGET samples. Each algorithm discovered optimal signal at various latent dimensionalities, but the top scoring features were generally identified in VAE and NMF models with large latent space dimensionalities (**Fig. 3e**). Although there were some potentially mischaracterized samples, feature 12 in VAE *k* = 50 robustly separated MYCN amplification status in NBL tumors (*t* = -18.5, *p* = 6.6 x 10^-38^) (**Fig. 3f**). This feature also distinguished MYCN amplification status in NBL cell lines [25] that were previously not used in training the compression model or for feature selection (*t* = -3.2, *p* = 4.2 x 10^-3^) (**Fig. 3g**).

### Assessing gene set coverage of compression models

We used gene sets from Molecular Signatures Database (MSigDB) and xCell [26–28] to interpret biological signals activated in compressed features across all latent dimensionalities, algorithms, and initializations. We applied a network projection approach to all compression algorithm weight matrices to determine gene set coverage. Briefly, we projected all compressed features onto a gene set network and assigned gene sets with the highest enrichment that passed an adjusted statistical significance threshold to each compressed feature (see methods for more details). We tracked coverage of three MSigDB gene set collections representing transcription factor (TF) targets, cancer modules, and Reactome pathways across latent dimensionalities in TCGA data (**Fig. 4**). In all cases, we observed higher gene set coverage in models with larger latent dimensionalities. Considering individual models, we observed high coverage in PCA, ICA, and NMF. In particular, ICA outperformed all other algorithms (**Fig. 4a**). However, while these methods showed the highest coverage, the features identified had relatively low enrichment scores compared to AE models (**Additional File 1: Figure S6**).

**Figure 4:**
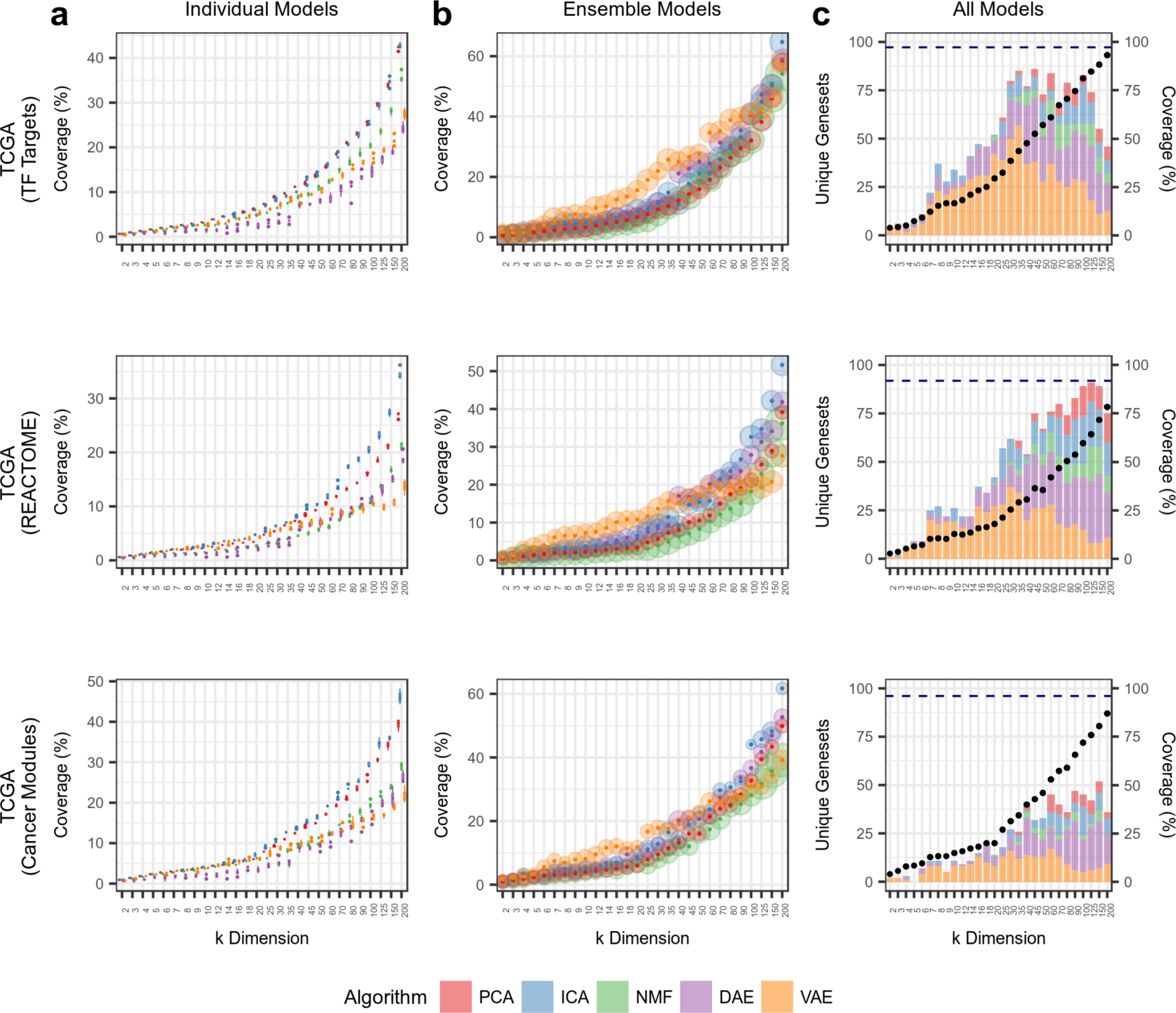
Assessing gene set coverage of specific gene set collections. Tracking results in TCGA data for three gene set collections representing transcription factor (TF) targets (C3TFT), Reactome pathways (C2CPREACTOME), and cancer modules (C4CM). **(a)** Tracking coverage in individual models, which represents the distribution of scores across five algorithm iterations. **(b)** Tracking coverage in ensemble models, which represents coverage after combining all five iterations into a single model. The size of the point represents relative enrichment strength. (**c**) Tracking coverage in all models combined within *k* dimensionalities. The number of algorithm-specific unique gene sets identified is shown as bar charts. Coverage for all models combined across all *k* dimensionalities is shown as a dotted navy blue line.

Aggregating all five random initializations into ensemble models, we observed substantial coverage increases, especially for AEs (**Fig. 4b**). VAE models had high coverage for all gene sets in intermediate dimensions, while DAE improved in higher dimensions. However, at the highest dimensions, ICA demonstrated the highest coverage. NMF consistently had the highest enrichment scores, but the lowest coverage (**Fig. 4b**). When considering all models combined (forming an ensemble of algorithm ensembles) within latent dimensionalities, we observed substantially increased coverage of all gene sets. However, most of the unique gene sets were contributed by the AE models (**Fig. 4c**).

Lastly, when we aggregated all BioBombe features across all algorithms and all latent dimensionalities together into a single model, we observed the highest gene set coverage (**Fig. 4c**). These patterns were consistent across other gene set collections and datasets (**Additional File 1: Figure S7**). In general, while models compressed with larger latent space dimensionalities had higher gene set coverage, many individual gene sets were captured with the highest enrichment in models with low and intermediate dimensionalities (**Additional File 1: Figure S8**). These results did not reveal a best method or dimensionality: Various biological representations are best discovered by using various compression algorithms with various latent space dimensionalities.

### Observing the latent dimensionality of specific tissue and cell type signatures

We measured the Pearson correlation between all samples’ gene expression input and reconstructed output. As expected, we observed increased mean correlation and decreased variance as the latent dimensionalities increased in TCGA data (**Fig. 5a**). We also observed similar patterns in GTEx and TARGET data (**Additional File 1: Figure S9**). Across all datasets, in randomly permuted data, we observed correlations near zero (**Additional File 1: Figure S9**). The correlation with real data was not consistent across all algorithms as PCA, ICA, and NMF generally outperformed the AE models.

**Figure 5:**
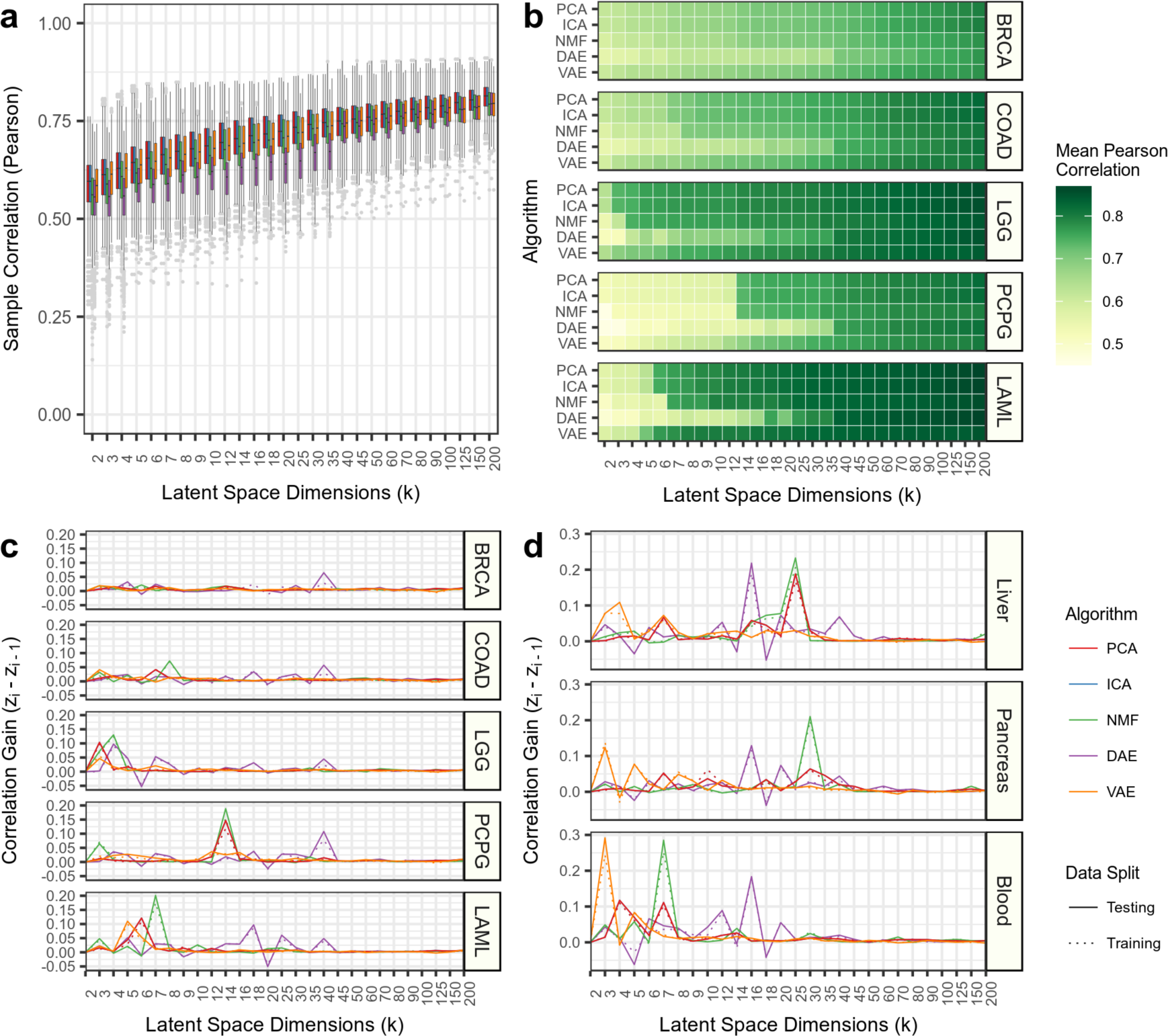
Different latent dimensionalities implicate different tissue types. **(a)** Sample Pearson correlation for all data in the testing data partition for The Cancer Genome Atlas (TCGA). The different algorithms follow the legend provided in panel d. **(b)** Mean Pearson correlation for select cancer types in the testing data partition. Pearson correlation gain between sequential latent dimensionalities for **(c)** select cancer types in TCGA and **(d)** select tissue-types in GTEx.

We tracked correlation differences across latent dimensionalities to determine the dimensionality at which specific sample types are initially detected. Most cancer types, including breast invasive carcinoma (BRCA) and colon adenocarcinoma (COAD), displayed relatively gradual increases in sample correlation as the latent dimensionality increased (**Fig. 5b**). However, in other cancer types, such as low grade glioma (LGG), pheochromocytoma and paraganglioma (PCPG), and acute myeloid leukemia (LAML), we observed large correlation gains with a single increase in latent dimensionality (**Fig. 5c**). We also observed similar performance spikes in GTEx data for several tissues including liver, pancreas, and blood (**Fig. 5d**). This sudden and rapid increase in correlation in specific tissues occurred at different latent dimensionalities for different algorithms, but was consistent across algorithm initializations.

To determine if this rapid increase was a result of models learning specific biological representations or if this observation represented a technical artifact, we more closely examined the sharp increase in GTEx blood tissue correlation between latent space dimensionalities 2 and 3 in VAE models (See **Fig. 5d**). We hypothesized that a difference in reconstruction for a specific tissue at such a low dimensionality could be driven by a change in the cell types captured by the model. We applied network projection of xCell gene sets to all compressed features in both VAE models. xCell gene sets represent computationally derived cell type signatures [27]. The top features identified for the VAE *k* = 2 model included skeletal muscle, keratinocyte, and neuronal gene sets (**Fig. 6a**). Skeletal muscle was the most significant gene set identified likely because it the tissue with the most samples in GTEx. Similar gene sets were enriched in the *k* = 3 model, but we also observed enrichment for a specific neutrophil gene set (“Neutrophils_HPCA_2”) (**Fig. 6a**). Neutrophils represent 50% of all blood cell types, which may explain the increased correlation in blood tissue observed in VAE *k* = 3 models. The features implicated using the network projection approach were similar to an overrepresentation analysis using high weight genes in both tails of the VAE *k* = 3 feature (**Additional File 1: Figure S10**).

**Figure 6:**
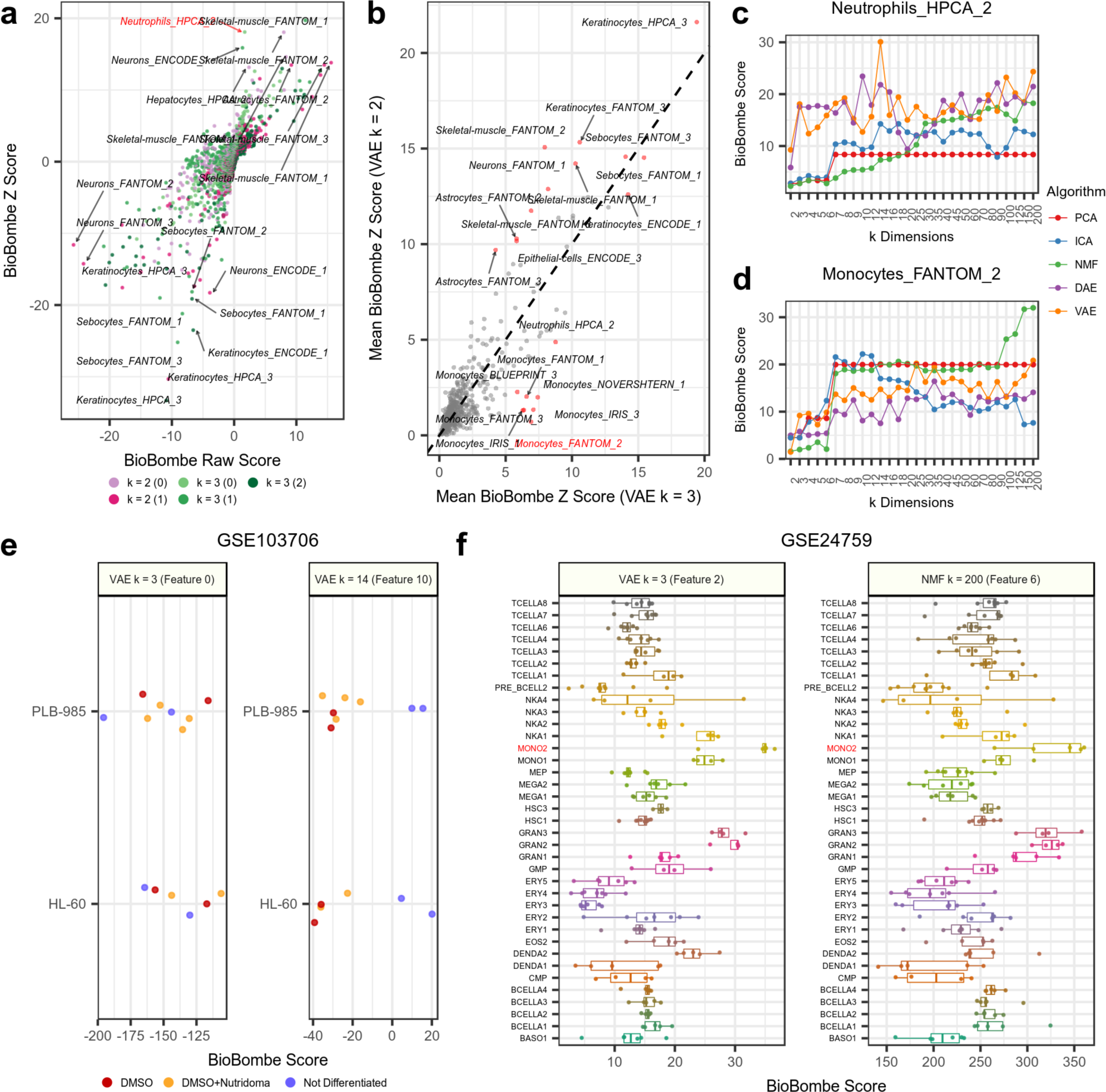
Interpreting blood cell types in GTEx using xCell gene sets. **(a)** Comparing BioBombe scores of all compressed latent features for variational autoencoder (VAE) models when bottleneck dimensionalities are set to *k* = 2 and *k* = 3. **(b)** Comparing mean BioBombe Z scores of aggregated latent features across two VAE models with *k* dimensionalities 2 and 3. Tracking the BioBombe Z scores of **(c)** “Neutrophils_HPCA_2” and **(d)** “Monocytes_FANTOM_2” gene sets across dimensionalities and algorithms. Only the top scoring feature per algorithm and dimensionality is shown. **(e)** Projecting the VAE feature *k* = 3 feature and the highest scoring feature (VAE *k* = 14) that best captures a neutrophil signature to an external dataset measuring neutrophil differentiation treatments (GSE103706). **(f)** Projecting the VAE *k* = 3 feature that best captures monocytes and the feature of the top scoring model (NMF *k* = 200) to an external dataset of isolated hematopoietic cell types (GSE24759).

We also calculated the mean absolute value z scores for xCell gene sets in all compression features for both VAE models with *k* = 2 and *k* = 3 dimensionalities (**Fig. 6b**). Again, we observed skeletal muscle, keratinocytes, and neuronal gene sets to be enriched in both models. However, we also observed a cluster of monocyte gene sets (including “Monocytes_FANTOM_2”) with enrichment in *k* = 3, but low enrichment in *k* = 2 (**Fig. 6b**). Monocytes are also important cell types found in blood, and it is probable these signatures also contributed to the increased correlation for the reconstructed blood samples in VAE *k* = 3 models. We provide the full list of xCell gene set genes for the neutrophil and monocyte gene sets that intersected with the GTEx data in **Additional File 3**.

We scanned all other algorithms and latent dimensionalities to identify other compression features with high enrichment scores in the “Neutrophils_HPCA_2” (**Fig. 6c**) and “Monocytes_FANTOM_2” gene sets (**Fig. 6d**). We observed stronger enrichment of the “Neutrophil_HPCA_2” gene set in AE models compared to PCA, ICA, and NMF, especially at lower latent dimensionalities. We observed the highest score for the “Neutrophil_HPCA_2” gene set at *k* = 14 in VAE models (**Fig. 6c**). The top VAE feature at *k* = 14 correlated strongly with the VAE feature learned at *k* = 3 (**Additional File 1: Figure S10**). Conversely, PCA, ICA, and NMF identified the “Monocytes_FANTOM_2” signature with higher enrichment than the AE models (**Fig. 6d**). We observed a performance spike at *k* = 7 for both PCA and NMF models, but the highest enrichment for “Monocytes_FANTOM_2” occurred at *k* = 200 in NMF models.

### Validating GTEx neutrophil and monocyte signatures in external datasets

We downloaded a processed gene expression dataset (GSE103706) that applied two treatments to induce neutrophil differentiation in two leukemia cell lines [29]. We hypothesized that projecting the dataset on the “Neutrophil_HPCA_2” signature would reveal differential scores in the treated cell lines. We observed large differences in sample activations of treated vs untreated cell lines in the top Neutrophil signature (VAE *k* = 14) (**Fig. 6e**). We also tested the “Monocytes_FANTOM_2” signature on a different publicly available dataset (GSE24759) measuring gene expression of isolated cell types undergoing hematopoiesis [30]. We observed increased scores for isolated monocyte cell population (MONO2) and relatively low scores for several other cell types for top VAE features (**Fig. 6f**).

We applied the top signatures for the neutrophil and monocyte gene sets to each external dataset (see **Fig. 6c, d**). We observed variable enrichment patterns across different algorithms and latent dimensionalities (**Additional File 1: Figure S11a**). These separation patterns were associated with network projection scores in NMF models, but were not consistent with other algorithms (**Additional File 1: Figure S11b**). Taken together, in this analysis we determined that 1) adding a single latent dimensionality that captured Neutrophil and Monocyte signatures improved signal detection in GTEx blood, 2) these gene expression signatures are enhanced at different latent dimensionalities and by different algorithms, and 3) these signatures generalized to external datasets that were not encountered during model training.

### Using BioBombe features in supervised learning applications

We used BioBombe compressed features in two supervised machine learning tasks. First, we trained logistic regression models using compressed BioBombe features from individual model iterations as input to predict each of the 33 different TCGA cancer types. Nearly all cancer types could be predicted with high precision and recall (**Additional File 1: Figure S12**). We observed multiple performance spikes at varying latent dimensionalities for different cancer types and algorithms, which typically occurred in small latent dimensionalities (**Fig. 7a**). Next, we input BioBombe features into the supervised classifier to predict samples with alterations in the top 50 most mutated genes in TCGA (**Additional File 1: Figure S13**). We focused on predicting four cancer genes and one negative control; *TP53*, *PTEN*, *PIK3CA*, *KRAS*, and *TTN* (**Fig. 7b**). *TTN* is a particularly large gene and is associated with a high passenger mutation burden and should provide no predictive signal [31]. As expected, we did not observe any signal in predicting *TTN* (**Fig. 7b**). Again, we observed performance increases at varying latent dimensionalities across algorithms. However, predictive signal for mutations occurred at higher latent dimensionalities compared to cancer types (**Fig. 7c**). Compared to features trained within algorithm and within iteration, an ensemble of five VAE models and an ensemble of five models representing one iteration of each algorithm (PCA, ICA, NMF, DAE, and VAE) identified cancer type and mutation status in earlier dimensionalities compared to single model iterations (**Fig 7c**). We also tracked the logistic regression coefficients assigned to each compression feature. DAE models consistently displayed sparse models, and the VAE ensemble and model ensemble also induced high sparsity (**Fig. 7d**).

**Figure 7:**
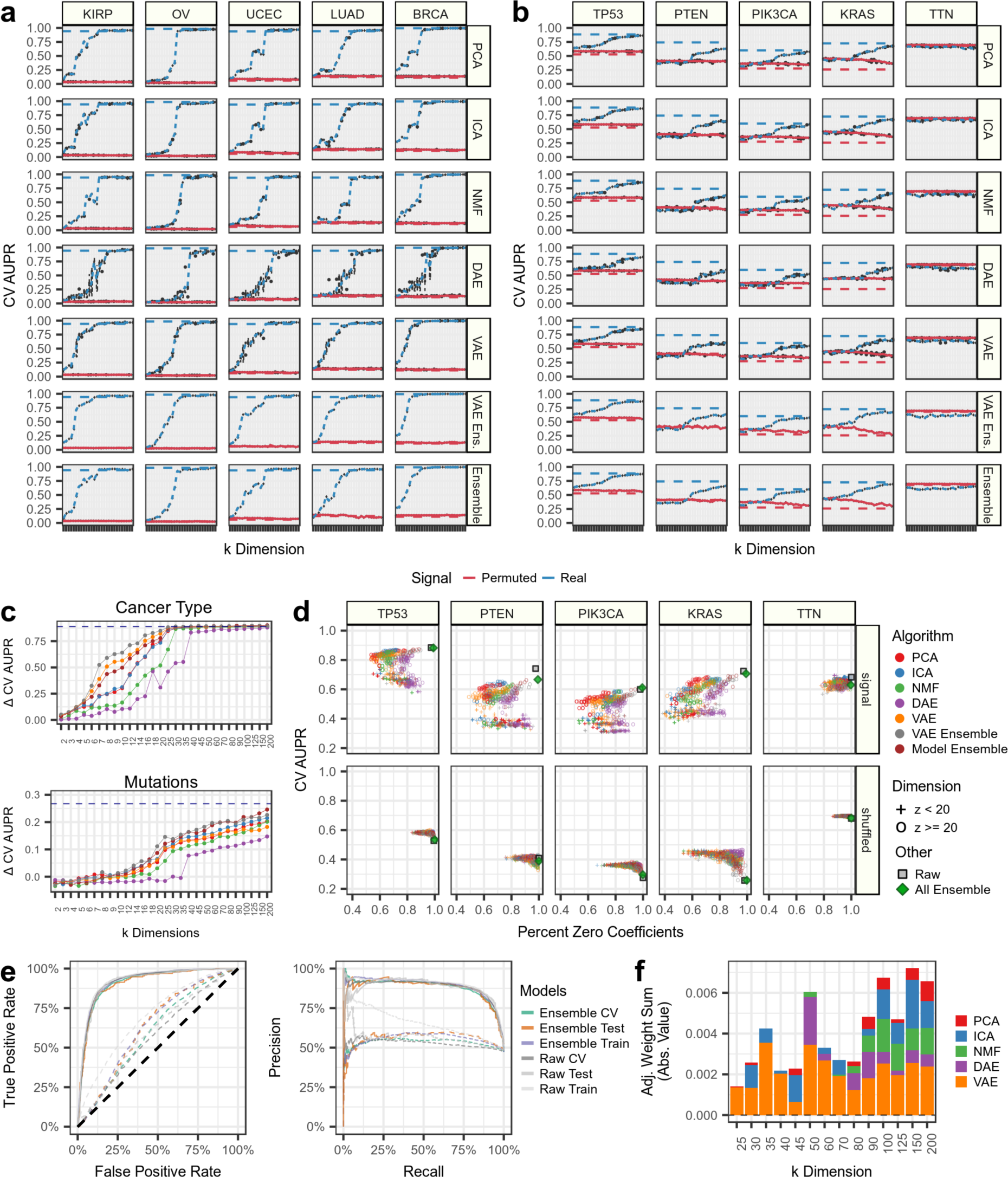
Using BioBombe sequential compression in The Cancer Genome Atlas (TCGA) as features in supervised machine learning tasks. Predicting. **(a)** cancer-type status and **(b)** gene mutation status for select cancer-types and important cancer genes using five compression algorithms and two ensemble models. The area under the precision recall (AUPR) curve for cross validation (CV) data partitions is shown. The blue lines represent predictions made with permuted data input into each compression algorithm. The dotted lines represent AUPR on untransformed RNAseq data. The dotted gray line represents a hypothetical random guess. **(c)** Tracking the average change in AUPR between real and permuted data across latent dimensionalities and compression models in predicting (*top*) cancer types and (*bottom*) mutation status. The average includes the five cancer types and mutations tracked in panels a and b. **(d)** Tracking the sparsity and performance of supervised models using BioBombe compressed features in real and permuted data. **(e)** Performance metrics for the all-compression feature ensemble model predicting *TP53* alterations. (*left*) Receiver operating characteristic (ROC) and (*right*) precision recall curves are shown. **(f)** The average absolute value weight per algorithm for the all-compression-feature ensemble model predicting *TP53* alterations. The adjusted scores are acquired by dividing by the number of latent dimensionalities in the given model.

Lastly, we trained logistic regression classifiers using all 30,850 BioBombe features generated across iterations, algorithms, and latent dimensionalities. These models were sparse and high performing; comparable to logistic regression models trained using raw features (**Fig. 7e**). Of all 30,850 compressed features in this model, only 317 were assigned non-zero weights (1.03%). We applied the network projection approach using Hallmark gene sets to interpret the biological signatures of the top supervised model coefficients. The top positive feature was derived from a VAE trained with *k* = 200. The top hallmarks of this feature included “ESTROGEN_RESPONSE_EARLY”, “ESTROGEN_RESPONSE_LATE”, and “P53_PATHWAY”. The top negative feature was derived from a VAE trained with *k* = 150 and was associated with hallmark gene sets including “BILE_ACID_METABOLISM”, “EPITHELIAL_MESENCHYMAL_TRANSITION”, and “FATTY_ACID_METABOLISM”. **Additional File 4** includes a full list of logistic regression coefficients and hallmark network projection scores. Overall, the features selected by the supervised classifier were distributed across algorithms and latent dimensionalities suggesting that combining signatures across dimensionalities and algorithms provided the best representation of the signal (**Fig. 7f**).

## Discussion

Our primary observation is that compressing complex gene expression data using multiple latent dimensionalities and algorithms improves biological representations. Across multiple latent dimensionalities, we identified optimal features to stratify sample sex, MYCN amplification, blood cell types, cancer types, and mutation status. These features generalized to other data, providing additional evidence for the intrinsic qualities of biological representations embedded in gene expression data [23,32–34]. Furthermore, the complexity of biological features was associated with the number of latent dimensionalities used. We predicted gene mutation using models with high dimensionality, but we detected cancer type with high accuracy using models with low dimensionality. In general, unsupervised learning algorithms applied to gene expression data extract biological and technical signals present in input samples. When applying these algorithms, researchers must determine how many latent dimensions to compress their input data into and different studies can have a variety of goals. For example, compression algorithms used for visualization can stratify sample groups based on the largest sources of variation [35–40]. In visualization settings, selecting a small number of latent dimensions is often best, and there is no need to compress data across multiple latent dimensionalities. However, if the analysis goal includes learning biological signatures to identify more subtle patterns in input samples, then there is not a single optimal latent dimensionality nor optimal algorithm. For example, though ICA and PCA represent rotations of each other, we found that the methods varied in their ability to capture specific biological signals into single features, which highlights the challenge of picking only a single algorithm. While compressing data into a single latent dimensionality will capture many biological signals, the “correct” dimensionality is not always clear, and several biological signatures may be better revealed by alternative latent dimensionalities.

If optimizing a single model, a researcher can use one or many criteria to select an appropriate latent dimensionality. Measurements such as Akaike information criterion (AIC), Bayesian information criterion (BIC), stability, and cross validation (CV) can be applied to a series of latent dimensionalities [41, 42]. Other algorithms, like Dirichlet processes, can naturally arrive at an appropriate dimensionality through several algorithm iterations [43]. Hidden layer dimensions of unsupervised neural networks are tunable hyperparameters defined by expected input data complexity and performance. However, applied to gene expression data, these metrics often provide conflicting results and unclear suggestions. In genomics applications, the method Thresher uses a combination of outlier detection and PCA to identify the optimal number of clusters [44]. Compression model stability can also be used to determine an optimal latent dimensionality in gene expression data [45]. By considering only reproducible features, ICA revealed 139 modules from nearly 100,000 publicly available gene expression profiles [46]. However, rather than using heuristics to select a biologically-appropriate latent dimensionality, a researcher may instead elect to compress gene expression data into many different latent space dimensionalities to generate many different feature representations.

There are many limitations in our evaluation of compressing gene expression data into many latent dimensionalities across multiple methods. First, our approach takes a long time to run. We are training many different algorithms across many different latent dimensionalities and iterations, which requires a lot of compute time (**Additional File 1: Figure S14**). However, because we are training many models independently, this task can be parallelized. Additionally, we did not evaluate dimensionalities above *k* = 200. It is likely that many more signatures can be learned, and possibly with even higher association strengths in higher dimensionalities for certain biology. We also did not focus on detecting compressed features that represent technical artifacts, which has already been widely explored [47, 48]. Moreover, we did not explore adding hidden layers in AE models. Many models trained on gene expression data have benefited from using multiple hidden layers in neural network architectures [7, 49]. Additional methods, like DeepLift, can be used to reveal gene importance values in internal representations of deep networks [50, 51].

An additional challenge is interpreting the biological content of the compressed gene expression features. Overrepresentation analysis (ORA) and gene set enrichment analysis (GSEA) are commonly applied but have significant limitations [26, 52]. ORA requires a user to select a cutoff, typically based on standard deviation, to build representative gene sets from each feature. ORA tests also do not consider the weights, or gene importance scores, in each compression feature. Conversely, GSEA operates on ranked features, but often requires many permutations to establish significance. Furthermore, ORA requires each tail of the compressed feature distribution to be interpreted separately in algorithms that also learn negative weights. The weight distribution is dependent on the specific compression algorithm, and the same cutoff may not be appropriate for all algorithms and all compressed features. Instead, we implemented a network projection based approach to quickly interpret compressed latent gene expression features [53, 54]. The approach is applied to the full and continuous distribution of gene weights, operates independently of the algorithm feature distribution, does not require arbitrary thresholds, and obviates the need to consider both tails of the distribution separately. Nevertheless, additional downstream experimental validation is necessary to determine if the constructed feature actually represents the biology it has been assigned.

## Conclusions

To enhance biological signature discovery, it is best to compress gene expression data using several algorithms and many different latent space dimensionalities. These signatures represent important biological signals, including various cell types, phenotypes, biomarkers, and other sample characteristics. We showed, through several experiments tracking lower dimensional gene expression representations, gene set coverage, and supervised learning performance, that optimal biological features are learned using a variety of latent space dimensionalities and different compression algorithms. As unsupervised machine learning continues to be applied to derive insight from biomedical datasets, researchers should shift focus away from optimizing a single model based on certain mathematical heuristics, and instead towards learning good and reproducible biological representations that generalize to alternative datasets regardless of compression algorithm and latent dimensionality.

## Methods

### Transcriptomic compendia acquisition and processing

We downloaded transcriptomic datasets from publicly available resources. We downloaded the batch-corrected TCGA PanCanAtlas RNAseq data from the National Cancer Institute Genomic Data Commons (https://gdc.cancer.gov/about-data/publications/pancanatlas). These data consisted of 11,069 samples with 20,531 measured genes quantified with RSEM and normalized with log transformation. We converted Hugo Symbol gene identifiers into Entrez gene identifiers and discarded non-protein coding genes and genes that failed to map. We also removed tumors that were measured from multiple sites. This resulted in a final TCGA PanCanAtlas gene expression matrix with 11,060 samples, which included 33 different cancer-types, and 16,148 genes. The breakdown of TCGA samples by cancer-type is provided in **Additional File 5**.

We downloaded the TPM normalized GTEx RNAseq data (version 7) from the GTEx data portal (https://gtexportal.org/home/datasets). There were 11,688 samples and 56,202 genes in this dataset. After selecting only protein-coding genes and converting Hugo Symbols to Entrez gene identifiers, we considered 18,356 genes. There are 53 different detailed tissue-types in this GTEx version. The tissues types included in these data are provided in **Additional File 5**.

Lastly, we retrieved the TARGET RNAseq gene expression data from the UCSC Xena data portal [55]. The TARGET data was processed through the FPKM UCSC Toil RNA-seq pipeline and was normalized with RSEM and log transformed [56]. The original matrix consists of 734 samples and 60,498 Ensembl gene identifiers. We converted the Ensembl gene identifiers to Entrez gene names and retained only protein-coding genes. This procedure resulted in a total of 18,753 genes measured in TARGET. There are 7 cancer-types profiled in TARGET and the specific breakdown is available in **Additional File 5**. All specific downloading and processing steps can be viewed and reproduced at https://github.com/greenelab/BioBombe/tree/master/0.expression-download.

### Training unsupervised neural networks

Autoencoders (AE) are unsupervised neural networks that learn through minimizing the reconstruction of input data after passing the data through one or several intermediate layers [57]. Typically, these layers are of a lower dimensionality than the input, so the algorithms must compress the input data. Denoising autoencoders (DAE) add noise to input layers during training to regularize solutions and improve generalizability [58]. Variational autoencoders (VAE) add regularization through an additional penalty term imposed on the objective function [59, 60]. In a VAE, the latent space dimensions (*k*) are penalized with a Kullback-Leibler (KL) divergence penalty restricting the distribution of samples in the latent space to Gaussian distributions. We independently optimized each AE model across a grid of hyperparameter combinations including 6 representative latent dimensionalities (described in **Additional File 2** and **Additional File 1: Figure S2**).

### Training compression algorithms across latent dimensionalities

Independently for each dataset (TCGA, GTEx, and TARGET), we performed the following procedure to train the compression algorithms. First, we randomly split data into 90% training and 10% testing partitions. We balanced each partition by cancer type or tissue type, which meant that each split contained relatively equal representation of tissues. Before input into the compression algorithms, we transformed the gene expression values by gene to the [0, 1] range by subtracting the minimum value and dividing by the range for each specific gene. We applied this transform independently for the testing and training partitions. We selected this range because it was compatible with all of the algorithms. We used the training set to train each compression algorithm. We used the scikit-learn implementations of PCA, ICA, and NMF, and the Tybalt implementations of VAE and DAE [8, 61].

After learning optimized compression models with the training data, we transformed the testing data using these models. We assessed performance metrics using both training and testing data to reduce bias. In addition to training with real data, we also trained all models with randomly permuted data. To permute the training data, we randomly shuffled the gene expression values for all genes independently. We also transformed testing partition data with models trained using randomly permuted data. Training with permuted data removes the correlational structure in the data and can help set performance metric baselines.

One of our goals was to assess differences in performance and biological signal detection across a range of latent dimensionalities (*k*). To this end, we trained all algorithms with various *k* dimensionalities including *k* = 2, 3, 4, 5, 6, 7, 8, 9, 10, 12, 14, 16, 18, 20, 25, 30, 35, 40, 45, 50, 60, 70, 80, 90, 100, 125, 150, and 200 for a total of 28 different dimensionalities. All of these models were trained independently. Lastly, for each *k* dimensionality we trained five different models initialized with five different random seeds. In total, considering the three datasets, five algorithms, randomly permuted training data, all 28 *k* dimensionalities, and five initializations, we trained 4,200 different compression models (**Additional File 2: Figure S1**). Therefore, in total, we generated 185,100 different compression features.

### Evaluating compression algorithm performance

We evaluated all compression algorithms on three main tasks: Reconstruction, sample correlation, and weight matrix stability. First, we evaluated how well the input data is reconstructed after passing through the bottleneck layer. Because the input data was transformed to a distribution between 0 and 1, we used binary cross entropy to measure the difference between algorithm input and output as a measure of reconstruction cost. The lower the reconstruction cost, the higher fidelity reconstruction, and therefore the higher proportion of signals captured in the latent space features. We also assessed the Pearson correlation of all samples comparing input to reconstructed output. This value is similar to reconstruction and can be quickly tracked at an individual sample level. Lastly, we used singular vector canonical correlation analysis (SVCCA) to determine model stability within and model similarity between algorithms and across latent dimensionalities [21]. The SVCCA method consisted of two distinct steps. First, singular value decomposition (SVD) was performed on two input weight matrices. The singular values that combined to reconstruct 98% of the signal in the data were retained. Next, the SVD transformed weight matrix was input into a canonical correlation analysis (CCA). CCA aligned different features in the weight matrix based on maximal correlation after learning a series of linear transformations. Taken together, SVCCA outputs a single metric comparing two input weight matrices that represents stability across model initializations and average similarity of two different models. Because we used the weight matrices, the similarity describes signature discovery. We use the distribution of SVCCA similarity measures across all pairwise algorithm initializations and latent dimensionalities to indicate model stability [21].

### Assessing gene expression signatures present in BioBombe features

We tested BioBombe sequentially compressed features to distinguish sample sex in GTEx and TCGA data, and MYCN amplification in TARGET NBL data. We tested all compression algorithms and latent space dimensionalities to determine the conditions in which these features were best captured. First, we selected tissue-types and cancer-types in the GTEx and TCGA sex analyses that were balanced by sex by selecting tissues with male to female ratios between 0.5 and 1.5. We performed a two-tailed independent t-test assuming unequal variance comparing male and female samples, and NBL samples with and without MYCN amplification. We applied the t-test to all compression features identified across algorithms, initializations, and dimensionalities. Shown in the figures are the top scoring feature per latent space dimensionality and algorithm.

We applied the optimal MYCN signature learned in TARGET to an alternative dataset consisting of a series of publicly available NBL cell lines [25]. The data were processed using STAR, and we accessed the processed FPKM matrix from figshare [62]. We transformed the dataset with the identified signatures using the following operation:

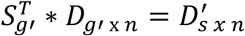

Where *D* represents the respective RNAseq data to transform, *S* represents the specific signature, *g’* represents the overlapping genes measured in both datasets, *n* represents samples, and *D’_s_* represents the signature scores in the transformed dataset. Of the 8,000 genes measured in TARGET data, 7,653 were also measured in external NBL cell line dataset (95.6%).

Using the sample activation scores for each of the top scoring features for sample sex in TCGA and GTEx, and MYCN amplification in TARGET and the validation set, we performed two tailed t-test with unequal variance comparing each group. For the TCGA and GTEx sex comparison, our t-test compared male vs. female activation scores. For the TARGET and NBL cell line analyses, our t-test compared MYCN amplified NBL samples vs. MYCN non-amplified NBL samples. We add t-test statistics and p values in each sub figure.

### Gene network construction and processing

We constructed networks using gene set collections compiled by version 6.2 of the Molecular Signatures Database (MSigDB) and cell types derived from xCell [26–28]. These gene sets represent a series of genes that are involved in specific biological processes and functions. We integrated all openly licensed MSigDB collections which included hallmark gene sets (H), positional gene sets (C1), curated gene sets (C2), motif gene sets (C3), computational gene sets (C4), Gene Ontology (GO) terms (C5), oncogenic gene sets (C6) and immunologic gene sets (C7). We omitted MSigDB gene sets that were not available under an open license (KEGG, BioCarta, and AAAS/STKE). The C2 gene set database was split into chemical and genetic perturbations (C2.CPG) and Reactome (C2.CP.Reactome). The C3 gene set was split into microRNA targets (C3.MIR) and transcription factor targets (C3.TFT). The C4 gene set was split into cancer gene neighborhoods (C4.CGN) and cancer modules (C4.CM). Lastly, the C5 gene set was split into GO Biological Processes (C5.BP), GO Cellular Components (C5.CC), and GO molecular functions (C5.MF). xCell represents a gene set compendia of 489 computationally derived gene signatures from 64 different human cell types. The number of gene sets in each curation is provided in **Additional File 6**. In BioBombe network projection, only a single collection is projected at a time.

To build the gene set network, we used hetnetpy [63]. Briefly, hetnetpy builds networks that include multiple node types and edge relationships. We used hetnetpy to build a single network containing all MSigDB collections and xCell gene sets listed above. The network consisted of 17,451 unique gene sets and 2,159,021 edges representing gene set membership among 20,703 unique gene nodes (**Additional File 6**). In addition to generating a single network using curated gene sets, we also used hetnetpy to generate 10 permuted networks. The networks are permuted using the XSwap algorithm, which randomizes connections while preserving node degree (i.e. the number of gene set relationships per gene) [64]. Therefore, the permuted networks are used to control for biases induced by uneven gene degree. We compared the observed score against the distribution of permuted network scores to interpret the biological signatures in each compression feature.

### Rapid interpretation of compressed gene expression data

Our goal was to quickly interpret the automatically generated compressed latent features learned by each unsupervised algorithm. To this end, we constructed gene set adjacency matrices with specific MSigDB or xCell gene set collections using hetnetpy software. We then performed the following matrix multiplication against a given compressed weight matrix to obtain a raw score for all gene sets for each latent feature.

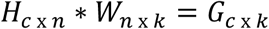

Where *H* represents the gene set adjacency matrix, *c* is the specific gene set collection, and *n* represents genes. *W* represents the specific compression algorithm weight matrix, which includes *n* genes and *k* latent space features. The output of this matrix multiplication, *G*, is represented by *c* gene sets and *k* latent dimensions. Through a single matrix multiplication, the matrix *G* tracks raw BioBombe scores.

Because certain hub genes are more likely to be implicated in gene sets and longer gene sets will receive higher raw scores, we compared *G* to the distribution of permuted scores against all 10 permuted networks.

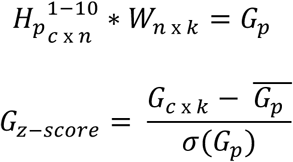

Where *H_P_^1-10^* represents the adjacency matrices for all 10 permuted networks and *G_p_* represents the distribution of scores for the same *k* features for all permutations. We calculated the z score for all gene sets by latent features (*G_z-score_*). This score represents the BioBombe Score. Other network-based gene set methods consider gene set influence based on network connectivity of gene set genes [53, 54]. Instead, we used the latent feature weights derived from unsupervised compression algorithms as input, and the compiled gene set networks to assign biological function.

We also compared the BioBombe network projection approach to overrepresentation analyses (ORA). We did not compare the approach to gene set enrichment analysis (GSEA) because evaluating single latent features required many permutations and did not scale to the many thousands of compressed features we examined. We implemented ORA analysis using a Fisher’s Exact test. The background genes used in the test included only the genes represented in the specific gene set collection.

### Calculating gene set coverage across BioBombe features

We were interested in determining the proportion of gene sets within gene set collections that were captured by the features derived from various compression algorithms. We considered a gene set “captured” by a compression feature if it had the highest positive or highest negative BioBombe z score compared to all other gene sets in that collection. We converted BioBombe z scores into p values using the pnorm() R function using a two-tailed test. We removed gene sets from consideration if their p values were not lower than a Bonferroni adjusted value determined by the total number of latent dimensionalities in the model.

We calculated coverage (C) by considering all unique top gene sets (*U*) identified by all features in the compression model (*w*) and dividing by the total number of gene sets in the collection (*T_C_*).

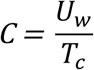

We calculated the coverage metric for all models independently (C*_i_*), for ensembles, or individual algorithms across all five iterations (*C_e_*), and for all models across *k* dimensions (*C_k_*). We also calculated the total coverage of all BioBombe features combined in a single model (C_all_). A larger coverage value indicated a model that captured a larger proportion of the signatures present in the given gene set collection.

### Downloading and processing publicly available expression data for neutrophil GTEx analysis

We used an external dataset to validate the neutrophil feature learned by compressing GTEx gene expression data into three latent dimensionalities. We observed that this feature contributed to improved reconstruction of blood tissue. To assess the performance of this neutrophil signature, we downloaded data from the Gene Expression Omnibus (GEO) with accession number GSE103706 [29]. RNA was captured in this dataset using Illumina NextSeq 500. The dataset measured the gene expression of several replicates of two neutrophil-like cell lines, HL-60 and PLB-985, which were originally derived from acute myeloid leukemia (AML) patients. The PLB-985 cell line was previously identified as a subclone of HL-60, so we expect similar signature activity between the two lines [65]. Gene expression of the two cell lines was measured with and without neutrophil differentiation treatments. Though DMSO is frequently used to solubilize compounds and act as an experimental control, it has been used to create neutrophil-like cells [66]. The validation dataset we used was generated to compare DMSO activity with untreated cells and cells treated with DMSO plus Nutridoma [29]. We tested the hypothesis that our neutrophil signature would distinguish the samples with and without neutrophil differentiation treatment. We transformed external datasets with the following operation:

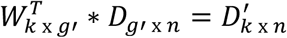

Where *D* represents the processed RNAseq data from GSE103706. Of 8,000 genes measured in *W*, 7,664 were also measured in *D* (95.8%). These 7,664 genes are represented by *g’*. All of the “Neutrophils_HPCA_2” signature genes were measured in *W*. *D’* represents the GSE103706 data transformed along the specific compression feature. Each sample in *D’* is then considered transformed by the specific signature captured in *k*. The specific genes representing “Neutrophils_HPCA_2” is provided in **Additional File 3**.

### Downloading and processing publicly available expression data for monocyte GTEx analysis

We used an additional external dataset to validate the identified monocyte signature. We accessed processed data for the publicly available GEO dataset with accession number GSE24759 [30]. The dataset was measured by Affymetrix HG-U133A (early access array) and consisted of 211 samples representing 38 distinct and purified populations of cells, including monocytes, undergoing various stages of hematopoiesis. The samples were purified from 4 to 7 independent donors each. Many xCell gene sets were computationally derived from this dataset as well [27]. Not all genes in the weight matrices were measured in the GSE24759 dataset. For this application, 4,645 genes (58.06%) corresponded with the genes used in the compression algorithms. Additionally, 168 out of 178 genes (94.38%) in the “Monocyte_FANTOM_2” gene set were measured (**Additional File 3**). We investigated the “Monocytes_FANTOM_2” signature because of its high enrichment in VAE *k* = 3 and low enrichment in VAE *k* = 2.

### Machine learning classification of cancer types and gene alterations in TCGA

We trained supervised learning classifiers using raw RNA-seq features and BioBombe-derived features. In general, we trained supervised machine learning models to predict cancer type from RNAseq features in TCGA PanCanAtlas RNAseq data. We implemented a logistic regression classifier with an elastic net penalty. The classifiers were controlled for mutation burden. More details about the specific implementation are described in Way et al. 2018 [67]. Here, we predicted all 33 cancer types using all 11,060 samples. These predictions were independent per cancer type, which meant that we trained models with the same input gene expression or BioBombe feature data, but used 33 different status matrices.

We also trained models to predict gene alteration status in the top 50 most mutated genes in the PanCanAtlas. These models were controlled for cancer type and mutation burden. We defined the status in this task using all non-silent mutations identified with a consensus mutation caller [68]. We also considered large copy number amplifications for oncogenes and deep copy number deletions for tumor suppressor genes as previously defined [69]. We used the threshold GISTIC2.0 calls for large copy amplifications (score = 2) and deep copy deletions (score = -2) in defining the status matrix [70]. For each gene alteration prediction, we removed samples with a hypermutator phenotype, defined by having log10 mutation counts greater than five standard deviations above the mean. For the mutation prediction task, we also did not include certain cancer types in training. We omitted cancer types if they had less than 5% or more than 95% representation of samples with the given gene alteration. The positive and negative sets must have also included at least 15 samples. We filtered out cancer types in this manner to prevent the classifiers from artificially detecting differences induced by unbalanced training sets.

We trained models with raw RNAseq data subset by the top 8,000 most variably expressed genes by median absolute deviation. The training data used was the same training set used for the BioBombe procedure. We also trained models using all BioBombe compression matrices for each latent dimension, and using real and permuted data. We combined compressed features together to form three different types of ensemble models. The first type grouped all five iterations of VAE models per latent dimensionality to make predictions. The second type grouped features of five different algorithms (PCA, ICA, NMF, DAE, VAE) of a single iteration together to make predictions. The third ensemble aggregated all features learned by all algorithms, all initializations, and across all latent dimensionalities, which included a total of 30,850 features. In total, considering the 33 cancer types, 50 mutations, 28 latent dimensionalities, ensemble models, raw RNAseq features, real and permuted data, and 5 initializations per compression, we trained and evaluated 32,868 different supervised models.

We optimized all of the models independently using 5-fold cross validation (CV). We searched over a grid of elastic net mixing and alpha hyperparameters. The elastic net mixing parameter represents the tradeoff between l1 and l2 penalties (where mixing = 0 represents an l2 penalty) and controls the sparsity of solutions [71]. Alpha is a penalty that tunes the impact of regularization, with higher values inducing higher penalties on gene coefficients. We searched over a grid for both hyperparameters (alpha = 0.1, 0.13, 0.15, 0.2, 0.25, 0.3 and mixing = 0.15, 0.16, 0.2, 0.25, 0.3, 0.4) and selected the combination with the highest CV AUROC. For each model, we tested performance using the original held out testing set that was also used to assess compression model performance.

### Evaluating model training time

We evaluated the execution time of training each compression algorithm for all three datasets across several latent dimensionalities. We used 8 representative latent dimensionalities: k = 2, 4, 10, 16, 25, 50, 80, and 200. We conducted the time analysis using a CPU machine with an Intel Core i3 dual core processer with 32 GB of DDR4 memory.

### Reproducible software

All code to perform all analyses and generate all results and figures is provided with an open source license at https://github.com/greenelab/biobombe [72]. All resources can be viewed and downloaded from https://greenelab.github.io/BioBombe/.

## Supporting information

neutrophil_monocyte_genesets

ensemble_coefficients_hallmarks_biobombe

cancer_tissue_types

metaedge_summary

## List of abbreviations

RNAseq: RNA sequencing
PCA: principal components analysis
ICA: independent components analysis
NMF: non-negative matrix factorization
AE: autoencoder
DAE: denoising autoencoder
VAE: variational autoencoder
TCGA: the cancer genome atlas
GTEx: genome tissue expression project
TARGET: therapeutically applicable research to generate effective treatments project
BRCA: breast invasive carcinoma
COAD: colon adenocarcinoma
LGG: low grade glioma
PCPG: pheochromocytoma and paraganglioma
LAML: acute myeloid leukemia
LUAD: lung adenocarcinoma
GEO: gene expression omnibus
ROC: receiver operating characteristic
PR: precision recall
AUROC: area under the receiver operating characteristic curve
AUPR: area under the precision recall curve
CV: cross validation
ORA: overrepresentation analysis
GSEA: gene set enrichment analysis
SVD: singular value decomposition
CCA: canonical correlation analysis
SVCCA: singular vector canonical correlation analysis
TF: transcription factor
DMSO: dimethyl sulfoxide

## Declarations

### Ethics approval and consent to participate

The TCGA, GTEx, and TARGET data used are publicly available and their use was previously approved by their respective ethics committees.

### Consent for publication

Not applicable.

### Availability of data and material

All data used and results generated in this manuscript are publicly available. All results of the analysis can be viewed and downloaded from https://greenelab.github.io/BioBombe/. The analyzed data can be accessed in the following locations: TCGA data can be accessed at https://gdc.cancer.gov/about-data/publications/pancanatlas, the GTEx data can be accessed at https://gtexportal.org/home/datasets, the TARGET data can be accessed at https://toil.xenahubs.net/download/target_RSEM_gene_fpkm.gz, the neutrophil validation data can be accessed using gene expression omnibus (GEO) accession number GSE103706 (https://www.ncbi.nlm.nih.gov/geo/query/acc.cgi?acc=GSE103706), the monocyte validation data can be accessed using GEO accession number GSE24759 (https://www.ncbi.nlm.nih.gov/geo/query/acc.cgi?acc=GSE24759). Software to reproduce the analyses, and all results generated in this manuscript can be accessed at https://github.com/greenelab/biobombe. The software has also been archived in an additional publicly available repository at https://zenodo.org/record/3460539.

Additionally, all BioBombe results are available in a series of versioned archives (TCGA BioBombe Results: https://zenodo.org/record/2110752); (GTEX BioBombe Results: https://zenodo.org/record/2300616); (TARGET BioBombe Results: https://zenodo.org/record/2222463); (Randomly permuted TCGA BioBombe Results: https://zenodo.org/record/2221216); (Randomly permuted GTEx BioBombe Results: https://zenodo.org/record/2386816); (Randomly Permuted TARGET BioBombe Results: https://zenodo.org/record/2222469). The full results of the TCGA classification analysis using BioBombe features is archived at https://zenodo.org/record/2535759.

### Authors’ contributions

GPW performed the analysis, wrote the BioBombe software, generated the figures, and wrote the manuscript. GPW and CSG designed the study and interpreted the results. MZ and DSH developed the network software. VR developed the website. All authors read, revised, and approved the final manuscript.

## Acknowledgements

We would like to thank Jaclyn Taroni, Yoson Park, and Alexandra Lee for insightful discussions and code review. We also thank Jo Lynne Rokita and John Maris for insightful discussions regarding the neuroblastoma analysis.

