## Supplementary Figures for "Sequential compression of gene expression across dimensionalities and methods reveals no single best method or dimensionality"

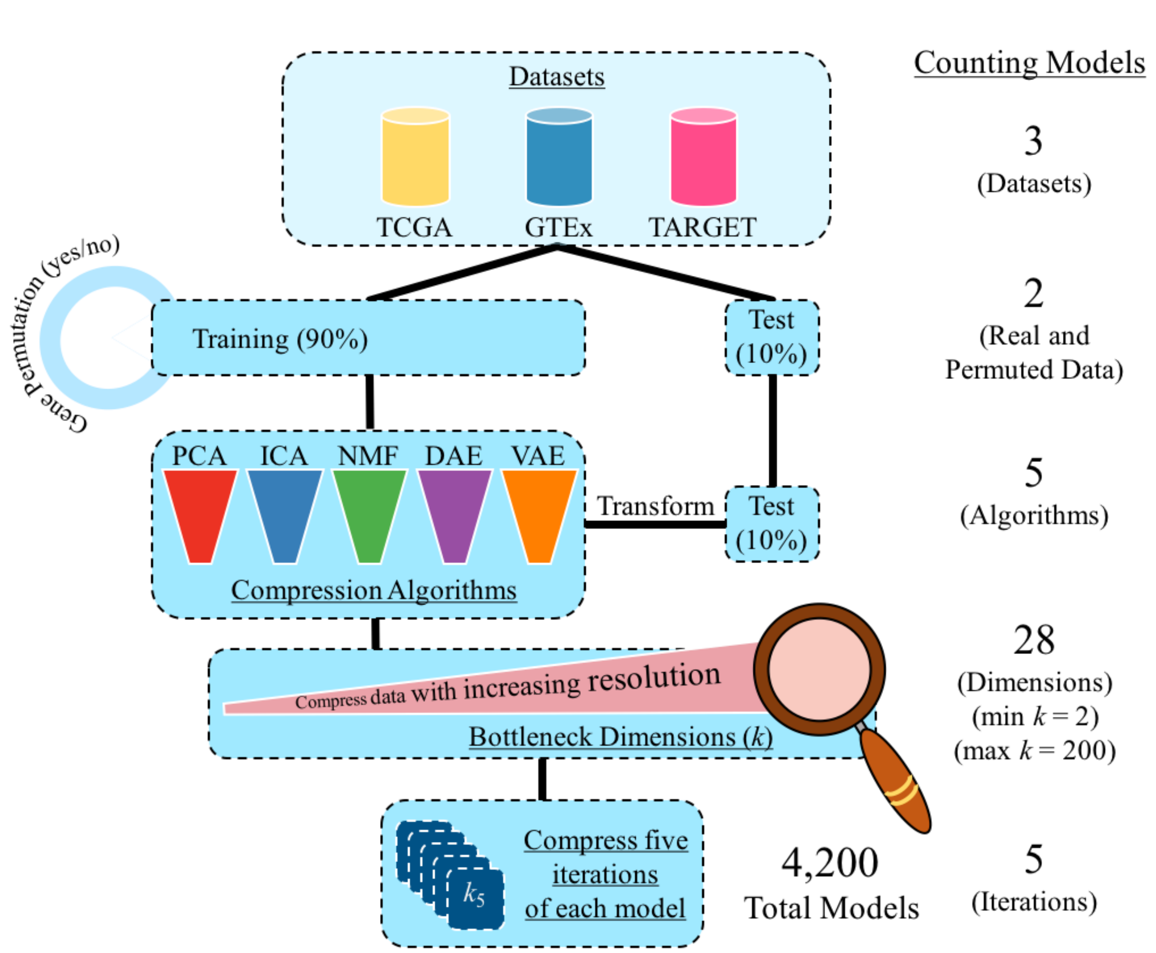

**Figure S1:** *Representing our BioBombe implementation workflow.* We independently apply our approach to three transcriptome datasets including The Cancer Genome Atlas PanCanAtlas Project (TCGA), Genome-Tissue Expression Project (GTEx), and Therapeutically Applicable Research to Generate Effective Treatments (TARGET) initiative. For each dataset, we split 90% of the data into a training data partition and 10% of the data into a testing data partition. The data is split to match the proportion of cancer types or tissue types in each partition. We also randomly permute the gene expression values by gene for all samples in the training set. We proceed with the downstream approach for both real and permuted data in parallel. We apply five compression algorithms including principal components analysis (PCA), independent components analysis (ICA), non-negative matrix factorization (NMF), denoising autoencoders (DAE), and variational autoencoders (VAE). We compress the testing data partition using the trained weights learned from the training set. We sequentially compress the input data into various latent dimensionalities ($k$) from 2 dimensions to 200 dimensions. We use $k$ = 2, 3, 4, 5, 6, 7, 8, 9, 10, 12, 14, 16, 18, 20, 25, 30, 35, 40, 45, 50, 60, 70, 80, 90, 100, 125, 150, and 200 for a total of 28 different dimensions. For each model, we train five independent times using five different random seed initializations. Combined, this yields a total of 4,200 different compression matrices.

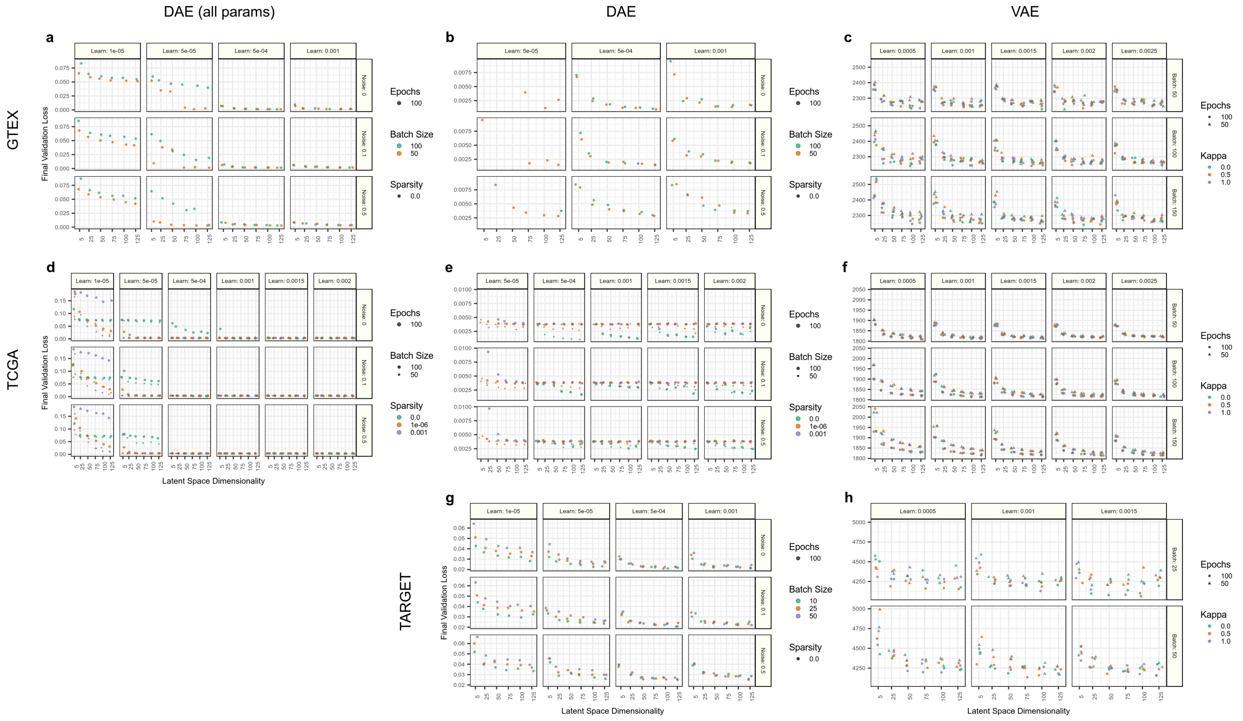

**Figure S2:** *Tracking performance across hyperparameter grid for autoencoder models.* We searched over a grid of various hyperparameter combinations for denoising autoencoder (DAE) and variational autoencoder (VAE) architectures. Training with several combinations of parameters in certain DAE models performed with high validation loss in Genome-Tissue Expression Project (GTEx) and The Cancer Genome Atlas PanCanAtlas Project (TCGA) data (panels **a** and **d**). We also show DAE performance with these particular hyperparameter combinations removed (panels **b** and **e**). Performance for all VAE hyperparameter combinations are shown (panels **c**, **f**, and **h**). Interestingly, no DAE model trained on Therapeutically Applicable Research to Generate Effective Treatments (TARGET) initiative data had very poor performance despite it being the smallest dataset (Panel **g**).

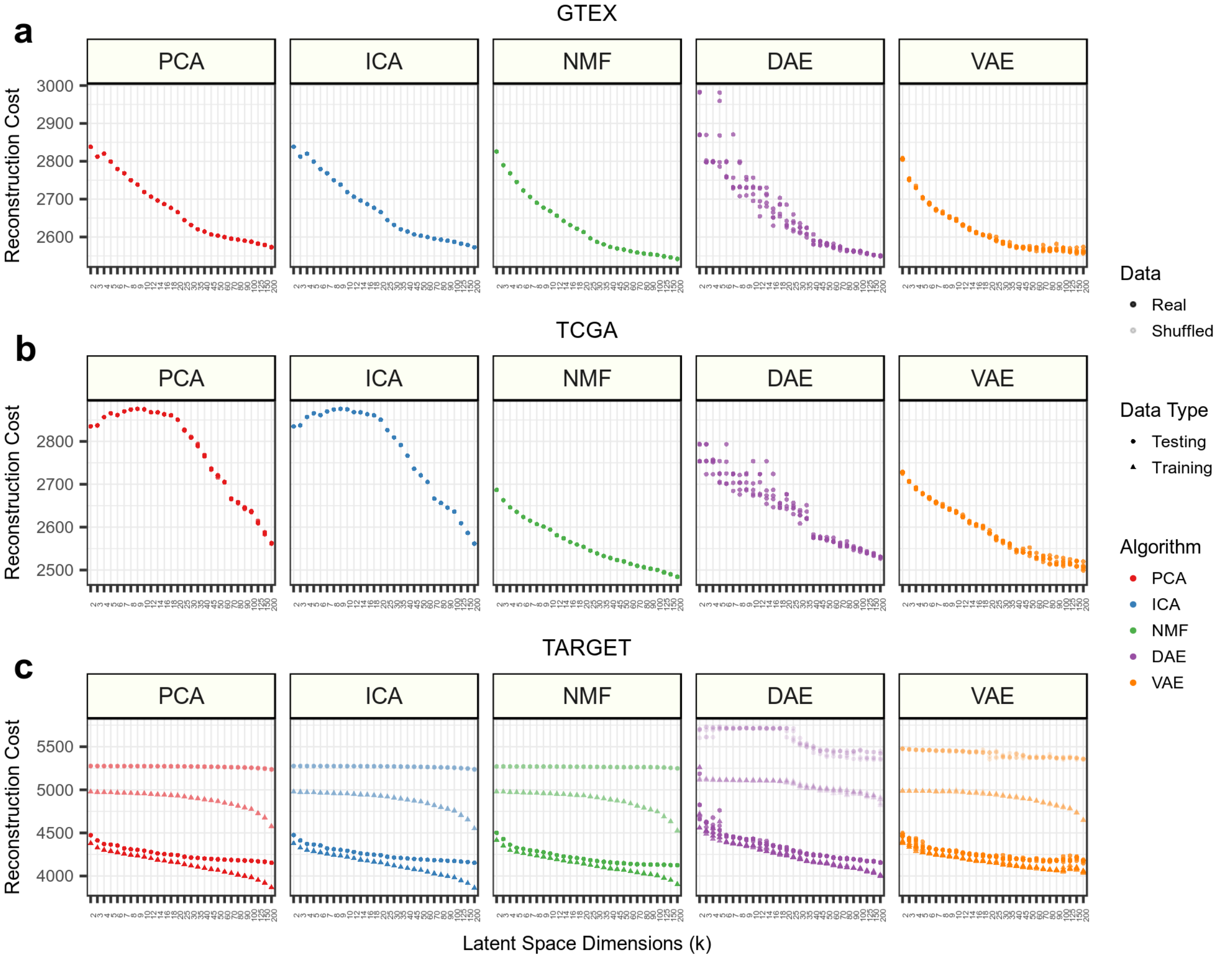

**Figure S3:** *Assessing reconstruction cost across datasets, algorithms, and latent dimensionalities.* Reconstruction performance for **(a)** Genome-Tissue Expression Project (GTEx) **(b)** The Cancer Genome Atlas PanCanAtlas Project (TCGA) and **(c)** Therapeutically Applicable Research to Generate Effective Treatments (TARGET) initiative data. Only real testing data is shown for GTEx and TCGA to highlight specific performance differences that would be unable to visualize with the other data present. Figures depicting all data are provided in our publicly available source code: https://github.com/greenelab/BioBombe/blob/master/4.analyze-components/

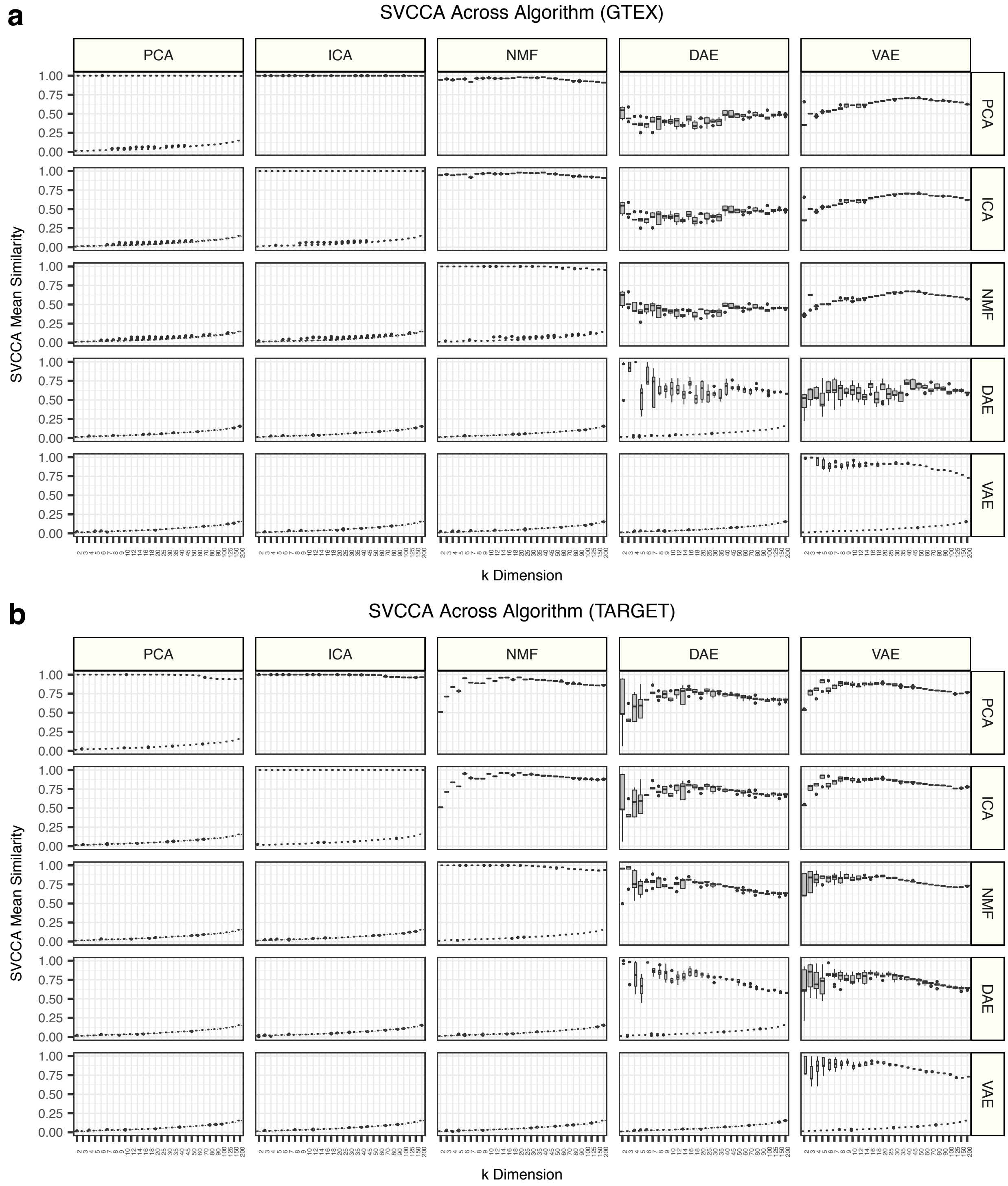

**Figure S4:** *Across algorithm stability as measured by singular vector canonical correlation analysis (SVCCA)*. Stability is measured for the weight matrices in **(a)** Genome-Tissue Expression Project (GTEx) and **(b)** Therapeutically Applicable Research to Generate Effective Treatments (TARGET). The boxplots represent all pairwise estimates of SVCCA mean similarity for all initializations (across seeds) for real data (upper triangle) and permuted data (lower triangle).

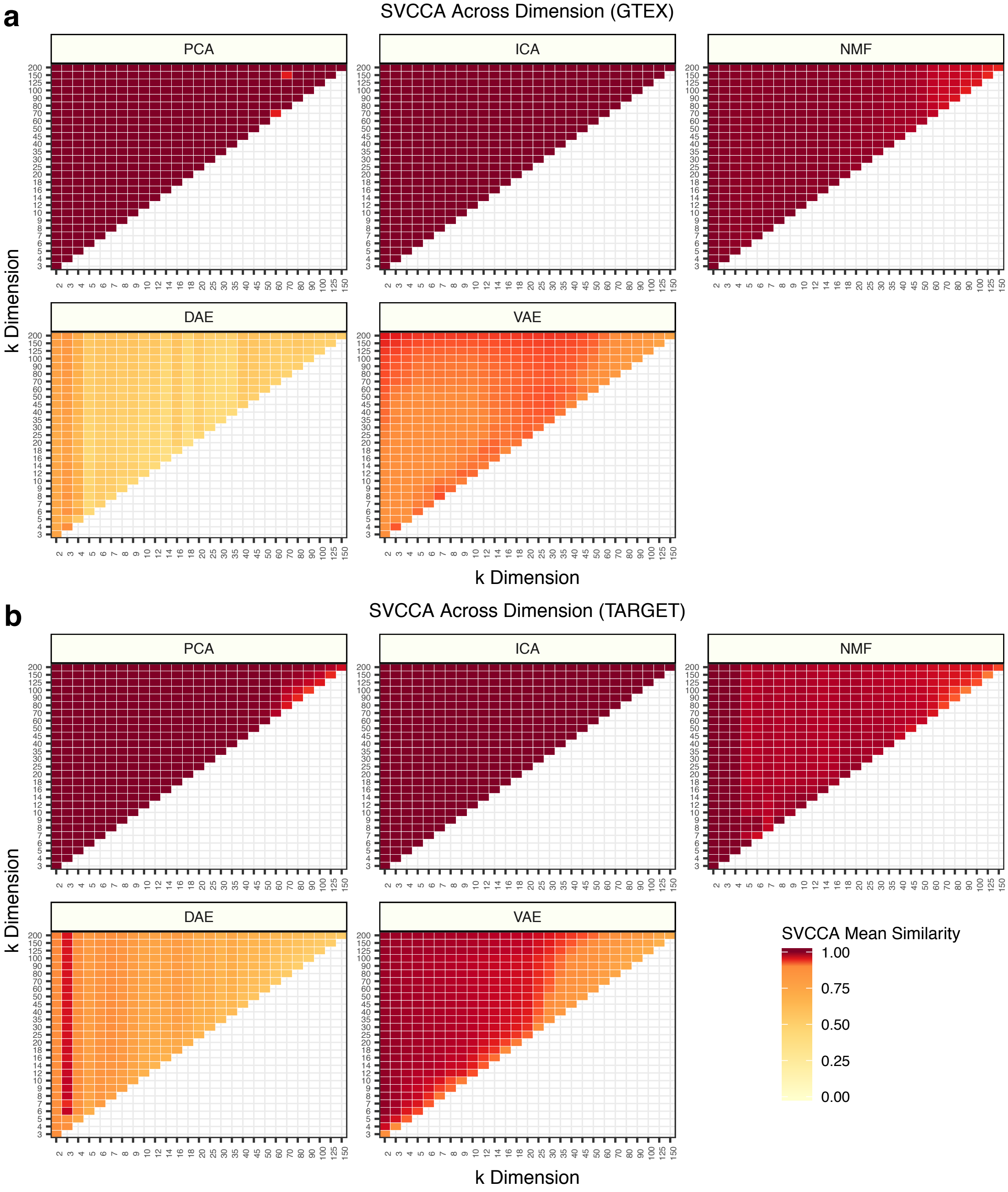

**Figure S5:** *Across latent dimensionalitystability as measured by singular vector canonical correlation analysis (SVCCA).* Stability is measured for the weight matrices in **(a)** Genome-Tissue Expression Project (GTEx) and **(b)** Therapeutically Applicable Research to Generate Effective Treatments (TARGET). SVCCA can be measured in two weight matrices of different dimensions. The mean similarity represents the mean of all pairwise estimates across all algorithm initializations. There is some numerical instability observed in the PCA assessment of both plots.

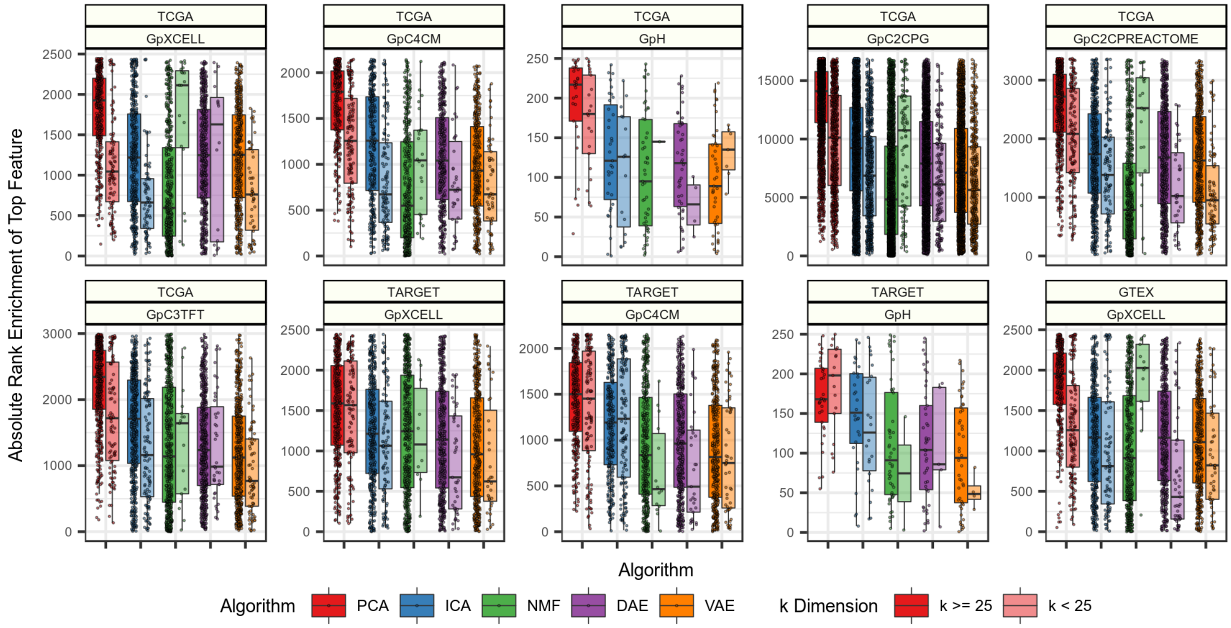

**Figure S6:** *Absolute ranking of top enriched gene set BioBombe z scores across algorithms.* We ranked all BioBombe z scores of top scoring gene sets within specific collections across algorithms. All gene sets within a specific collection are visualized within each algorithm box plot and whether or not they were identified as a top feature in model dimensionalitiesless than $k$ = 25.

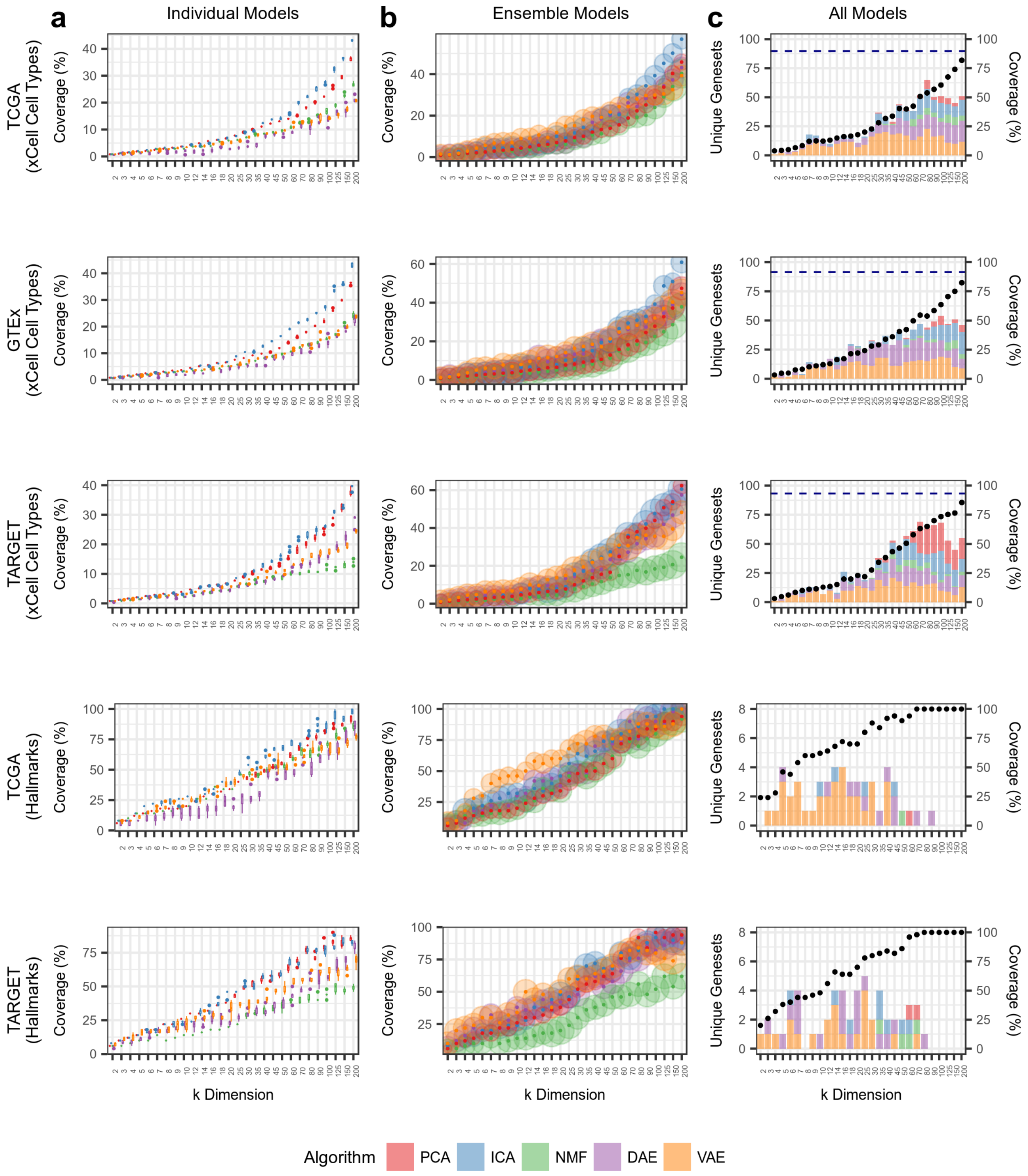

**Figure S7:** *Assessing gene set coverage of gene set collections and datasets.* Tracking results in TCGA data for two gene set collections representing cell types (xCell) and hallmark pathways (H), results in GTEx data for cell types, and results in TARGET data for cell types and hallmarks. Coverage represents the percentage of gene sets represented as the top feature. (**a**) Coverage in individual models, which represents the score distribution across five algorithm iterations. (**b**) Coverage in ensemble models, which combines all five iterations in a single model. (**c**) Coverage in all models within *k* dimensions. The number of algorithm-specific unique gene sets identified is shown as bar charts. Coverage for every model combined is shown as a dotted navy blue line.

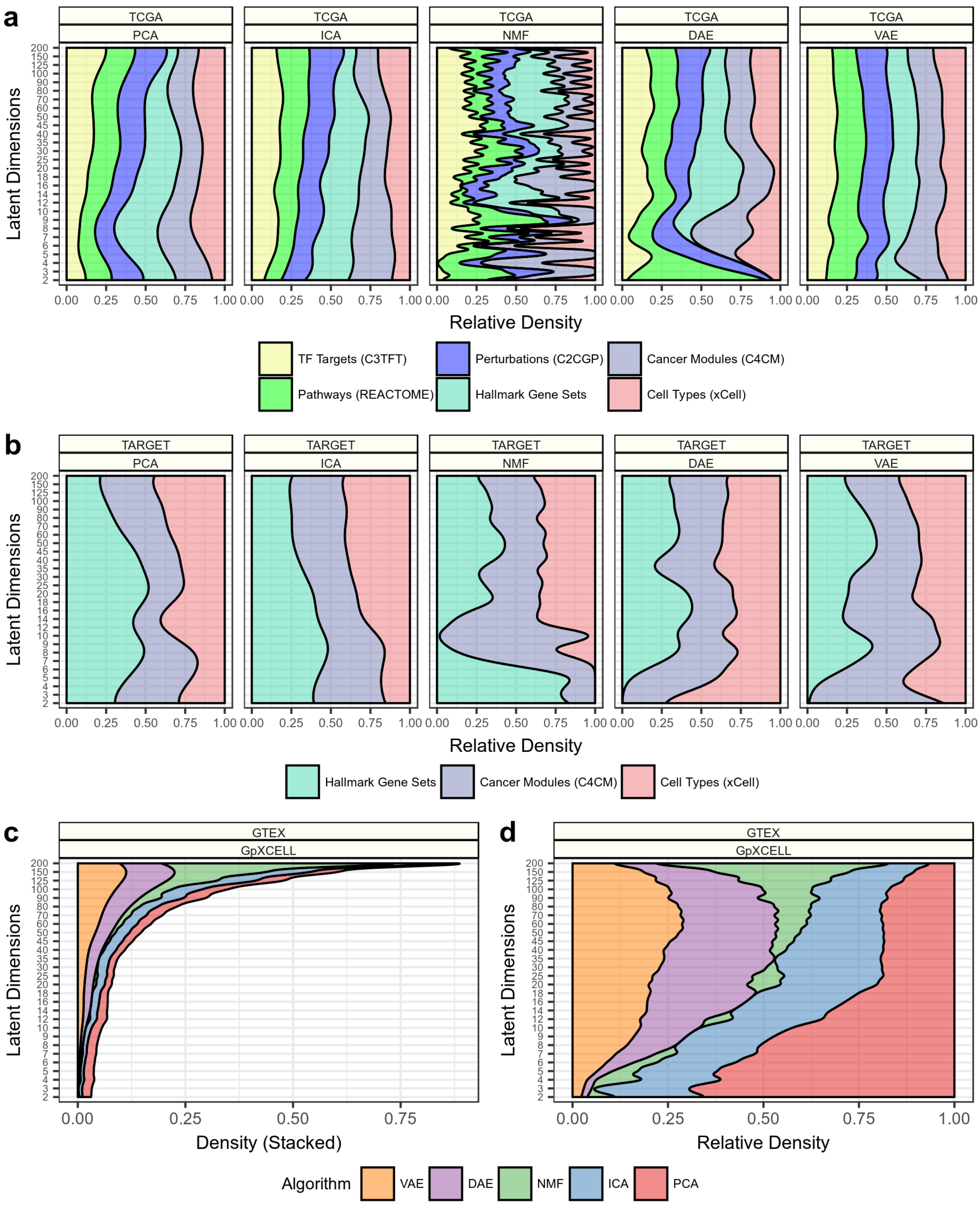

**Figure S8:** *Tracking the dimensionalities of highest BioBombe enrichment signal for gene sets, algorithms, and datasets.* The latent space dimension at which a gene set was identified with the highest enrichment across latent dimensionalities is shown. Observing the relative density of top features identified for several gene set collections across algorithms in (**a**) TCGA (**b**) TARGET data. Comparing (**c**) total counts and (**d**) relative density of xCell gene sets enrichment across latent dimensionality in GTEx data.

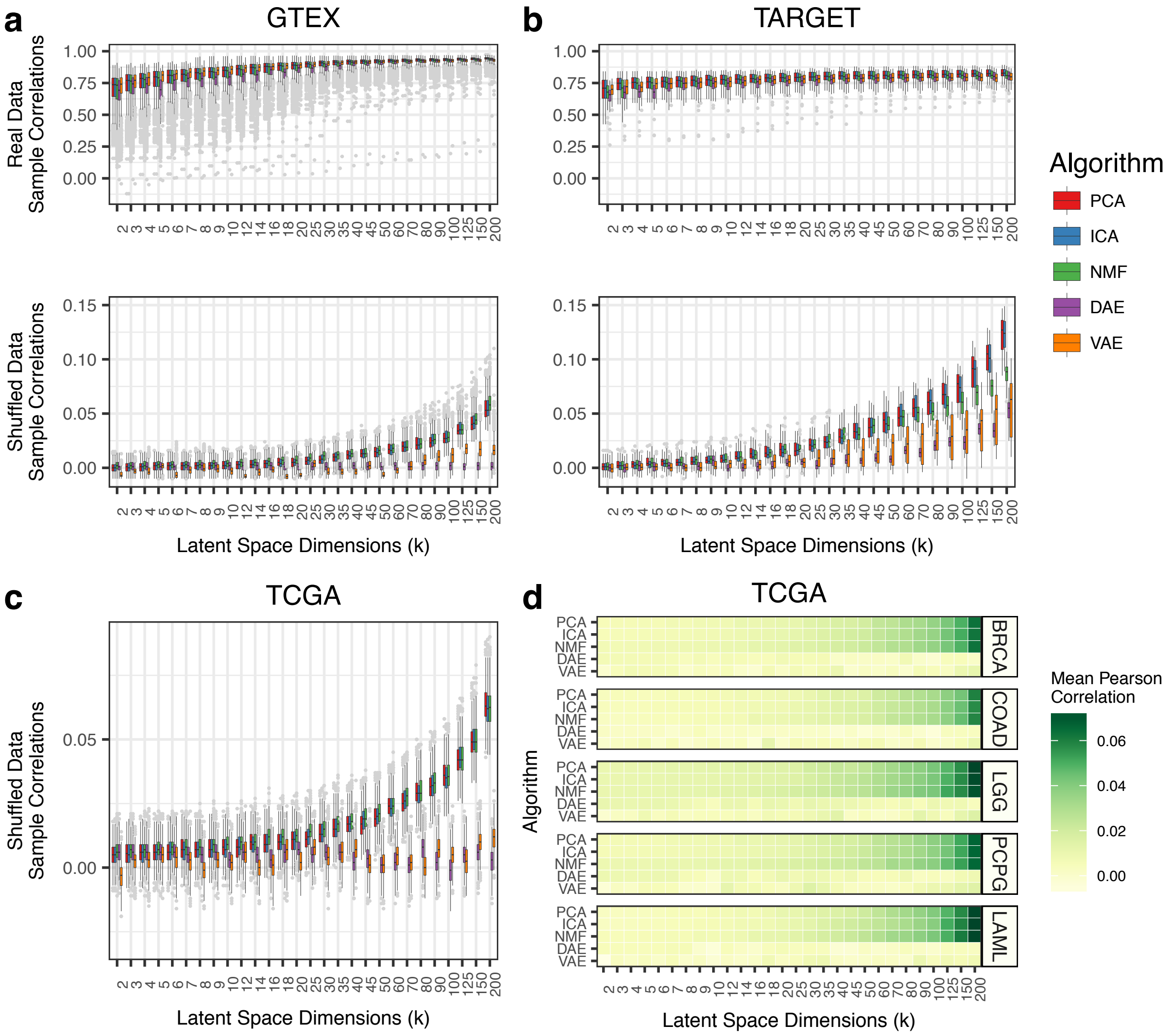

**Figure S9:** *Pearson correlation between input and reconstructed samples in real and permuted data.* Pearson sample correlation for real (top) and permuted (bottom) data in **(a)** Genome-Tissue Expression Project (GTEx) and **(b)** Therapeutically Applicable Research to Generate Effective Treatments (TARGET). **(c)** Pearson correlations in permuted data from The Cancer Genome Atlas PanCanAtlas Project (TCGA). **(d)** Pearson correlations between input and reconstructed output in permuted data to mirror select cancer-types in Figure 3. The data are permuted before input to the compression algorithms. Results across all specific cancer-types and tissue-types for GTEx, TARGET, and TCGA are provided in: https://github.com/greenelab/BioBombe/blob/master/4.analyze-components/

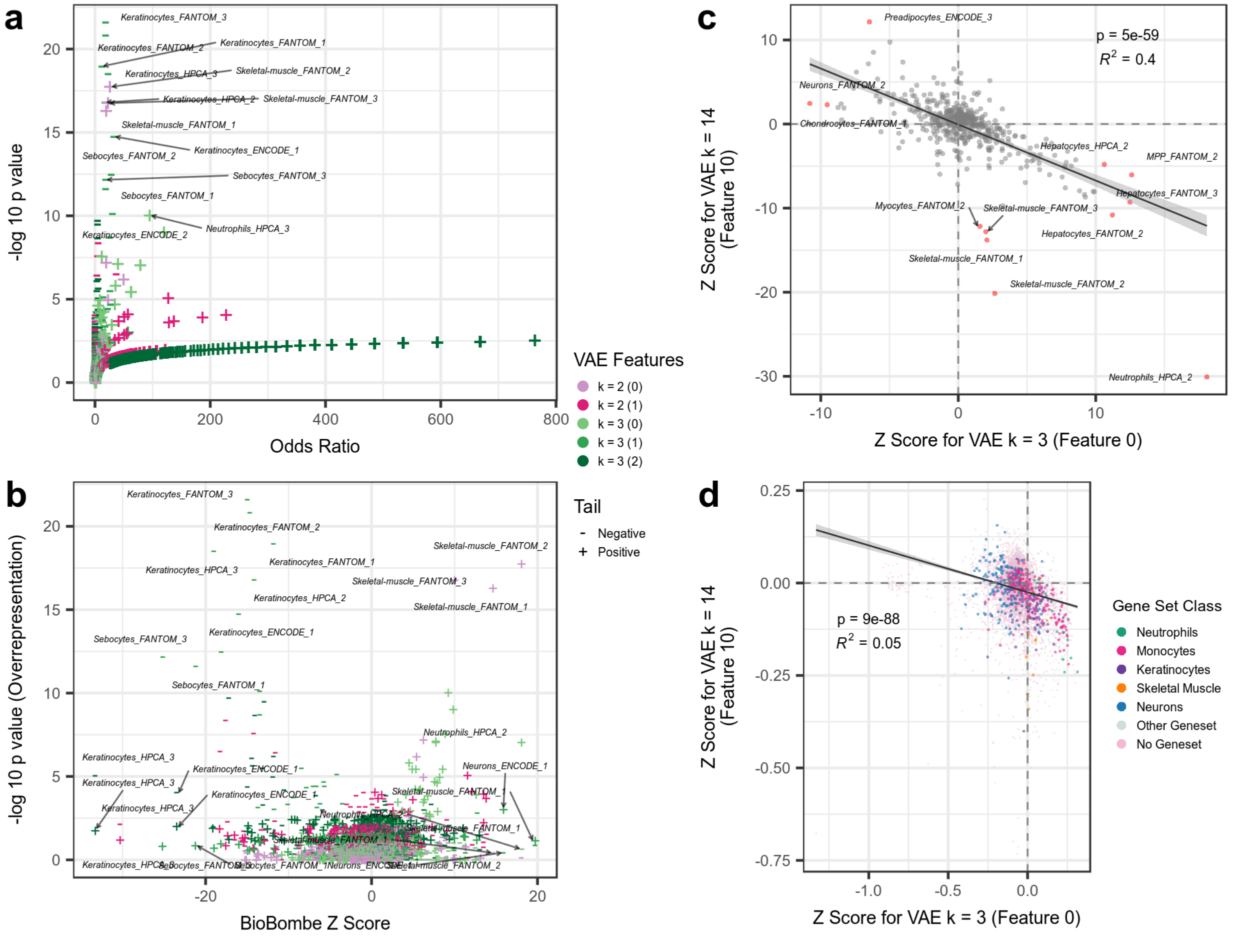

**Figure S10:** *Comparing overrepresentation analysis (ORA) with BioBombe derived scores for variational autoencoder (VAE) model for k = 3.* VAE trained on gene expression data from Genome-Tissue Expression Project (GTEx). Enrichment results are shown for the xCell geneset collection. **(a)** ORA analysis applied to feature 0 for VAE *k* = 3. **(b)** Comparing BioBombe Z score against ORA significance for the same VAE feature. The two tails of the feature distribution are shown for the ORA analysis and were extracted from the high weight genes. High weight genes were defined as having higher or lower weights than 2 standard deviations of the mean weight distribution. Comparing the VAE *k* = 3 feature that captures the neutrophil gene set to the VAE *k* = 14 feature that captures this gene set with the highest enrichment for **(c)** xCell gene set BioBombe Z scores and **(d)** compression algorithm gene weights.

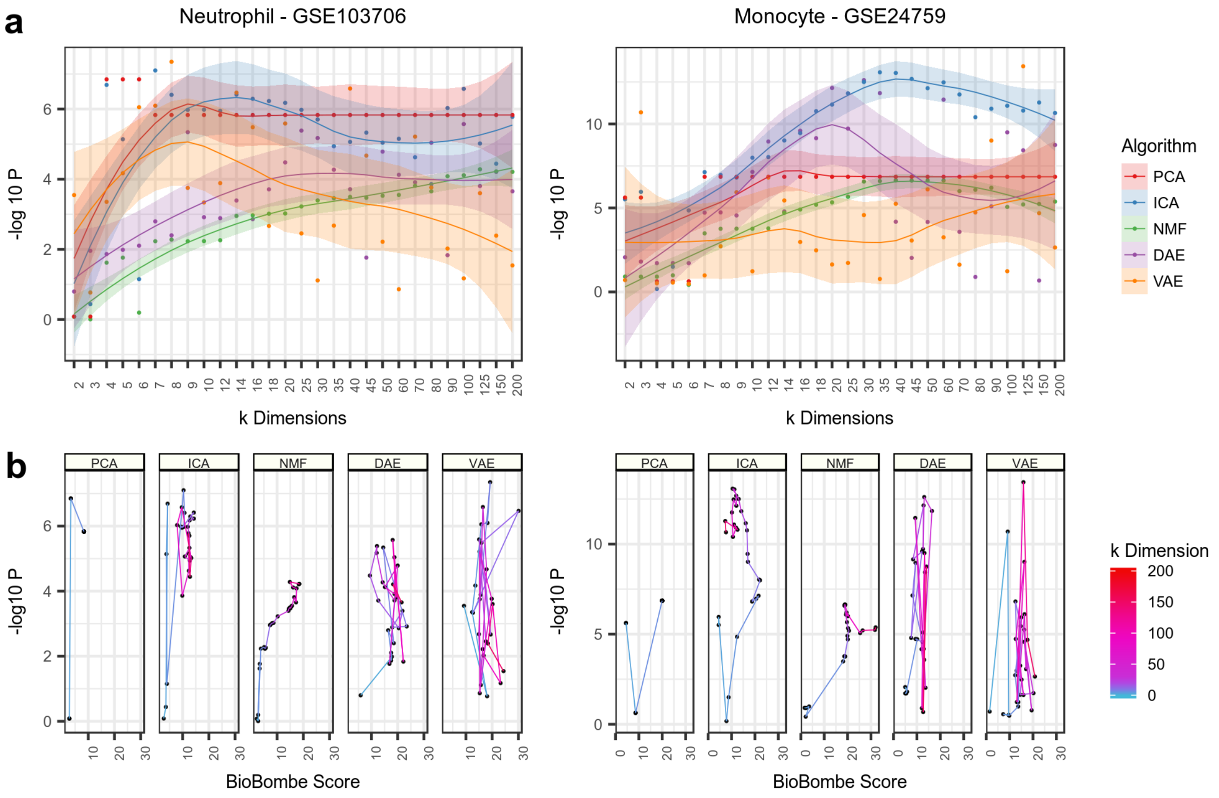

**Figure S11:** *Applying optimal features implicating Neutrophil and Monocyte specific signatures to two external datasets.* **(a)** Visualizing the highest enriched feature across algorithms and latent dimensionalities based on the –log 10 p value of an independent t-test comparing neutrophil (*left*) and monocyte (*right*) publicly available data after signature transformation. **(b)** Tracking the relationship between t-test significance and BioBombe scores of top gene sets across latent space dimensionalities. The increasing color intensity represents increasing dimension.

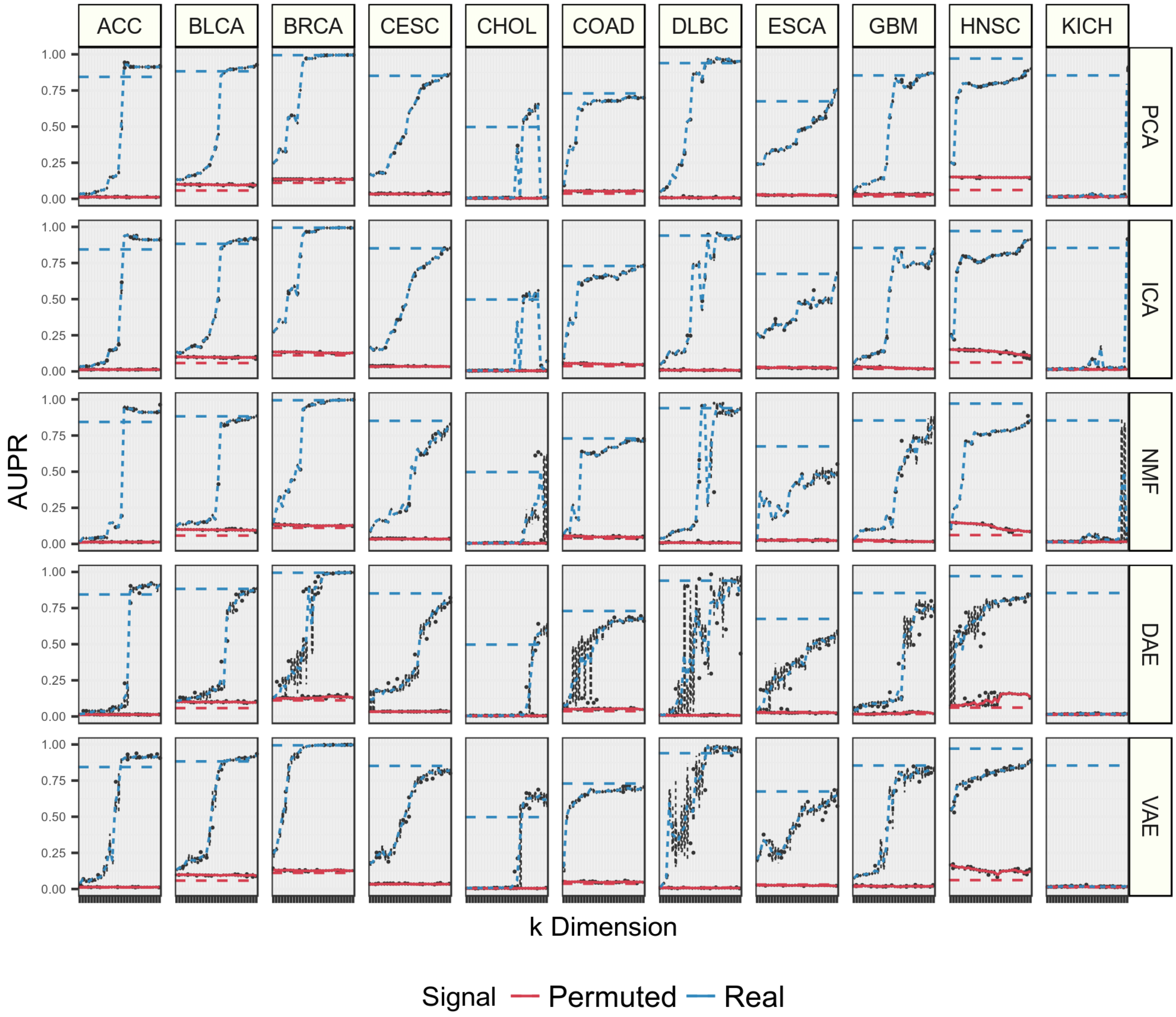
ACC
BLCA
BRCA
CESC
CHOL
COAD
DLBC
ESCA
GBM
HNSC
KICH
PCA
ICA
NMF
DAE
VAE
AUPR
1.00
0.75
0.50
0.25
0.00
k Dimension
Signal
Permuted
Real

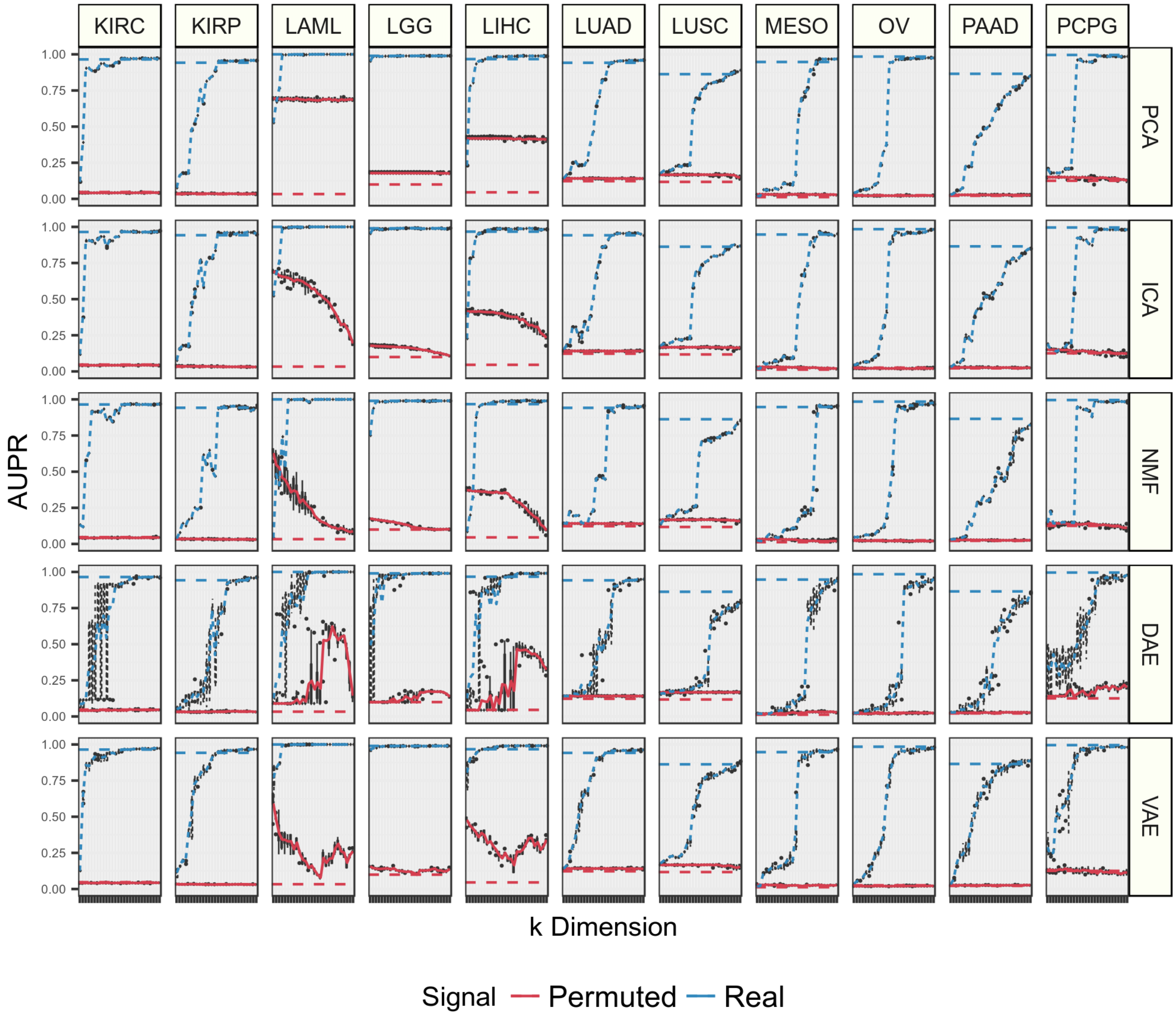
KIRC
KIRP
LAML
LGG
LIHC
LUAD
LUSC
MESO
OV
PAAD
PCPG
PCA
ICA
NMF
DAE
VAE
AUPR
1.00
0.75
0.50
0.25
0.00
k Dimension
Signal
Permuted
Real

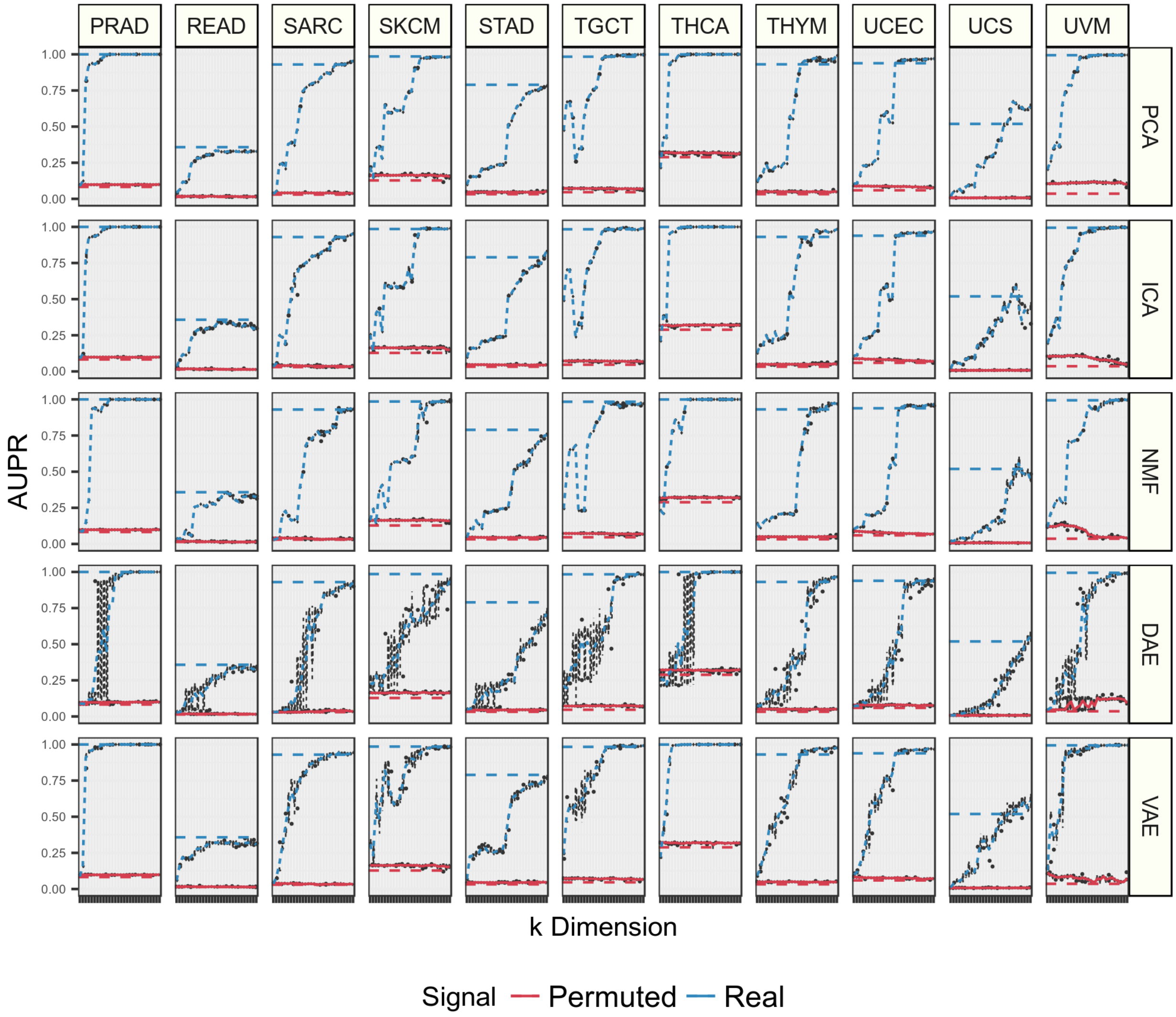

**Figure S12:** *Predicting 33 cancer-types in The Cancer Genome Atlas (TCGA) PanCanAtlas Project with features derived from different compression algorithms across latent dimensionalities.* We optimized logistic regression classifiers using compressed features for real and permuted data derived from five compression algorithms across dimensions. All 33 cancer-types are split across a series of three figures and are presented in alphabetical order. The area under precision recall (AUPR) for training data in cross validation intervals are shown. The blue lines represent compressed features derived from real training data input into the compression algorithms. The red lines represent compressed permuted data. The dotted lines in red and blue indicate performance with the raw RNAseq features and serve as baselines. All models, including the permuted data models, include covariates for cancer-type and log 10 mutation burden. The distribution of performance is provided in boxplots for all five algorithm initializations. The lines connect the mean of these algorithm initializations.

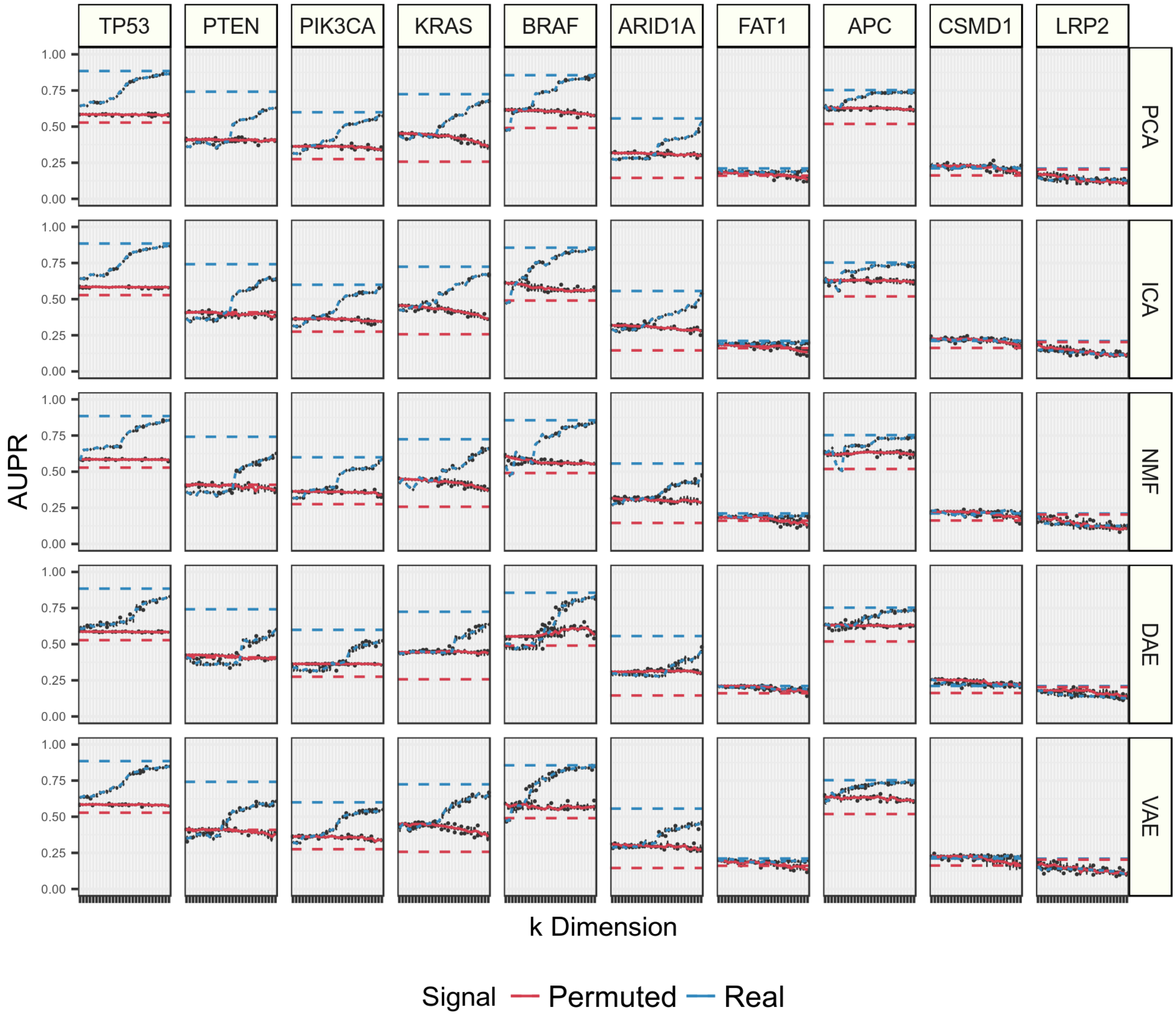
TP53
PTEN
PIK3CA
KRAS
BRAF
ARID1A
FAT1
APC
CSMD1
LRP2
PCA
ICA
NMF
DAE
VAE
AUPR
1.00
0.75
0.50
0.25
0.00
k Dimension
Signal
Permuted
Real

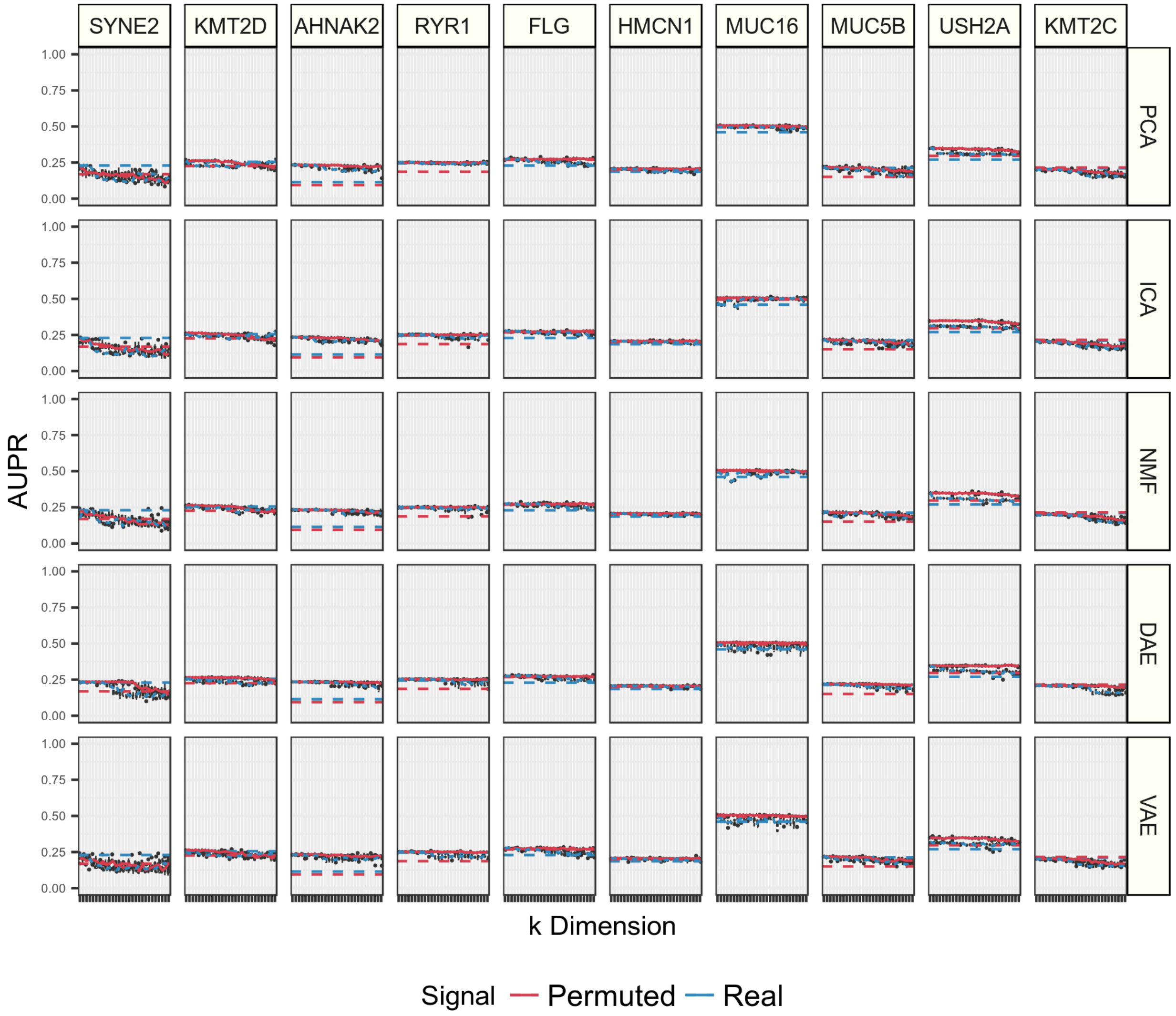

SYNE2
KMT2D
AHNAK2
RYR1
FLG
HMCN1
MUC16
MUC5B
USH2A
KMT2C
PCA
ICA
NMF
DAE
VAE
AUPR
1.00
0.75
0.50
0.25
0.00
k Dimension
Signal
Permuted
Real

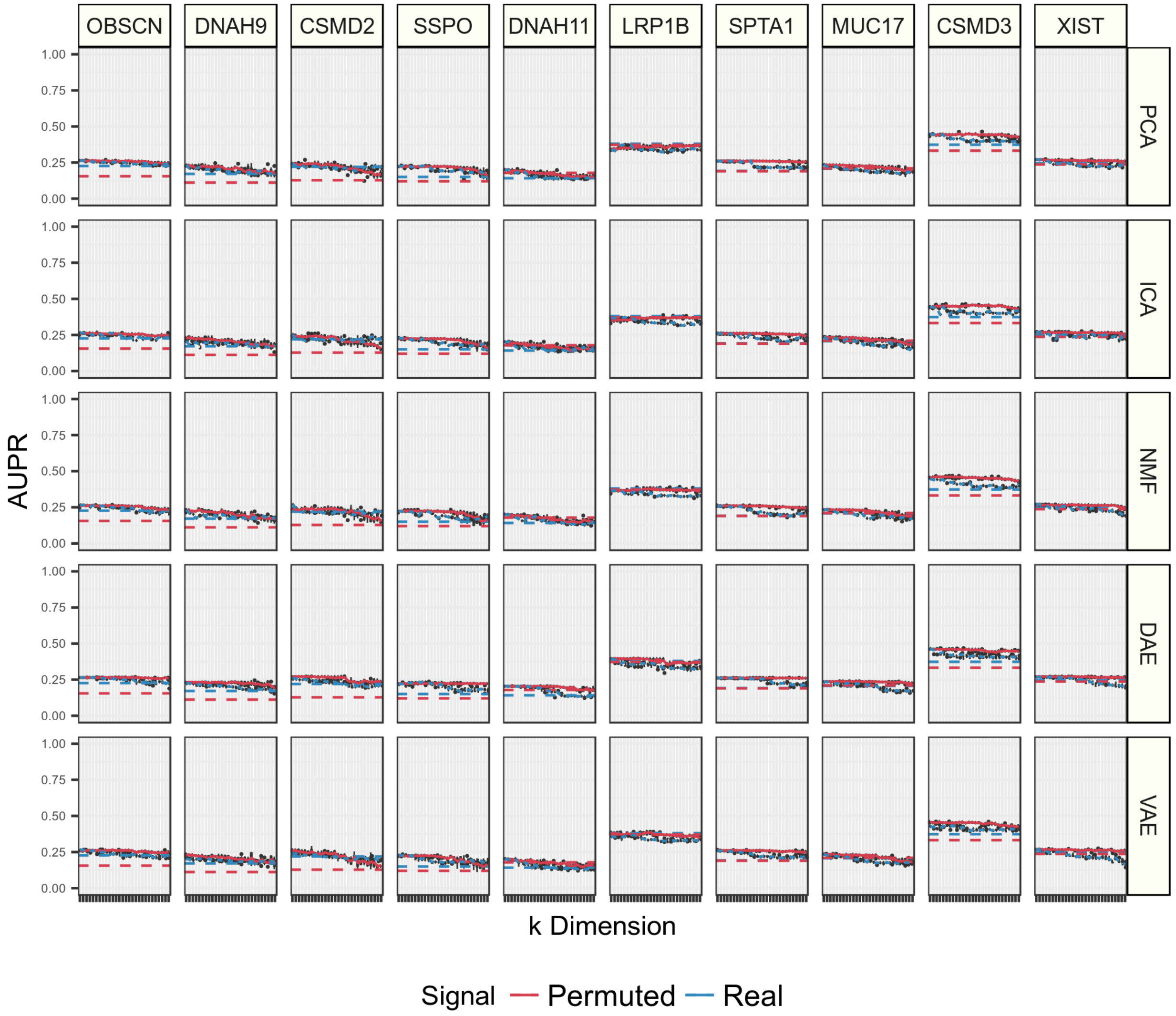
OBSCN
DNAH9
CSMD2
SSPO
DNAH11
LRP1B
SPTA1
MUC17
CSMD3
XIST
PCA
ICA
NMF
DAE
VAE
AUPR
1.00
0.75
0.50
0.25
0.00
k Dimension
Signal
Permuted
Real

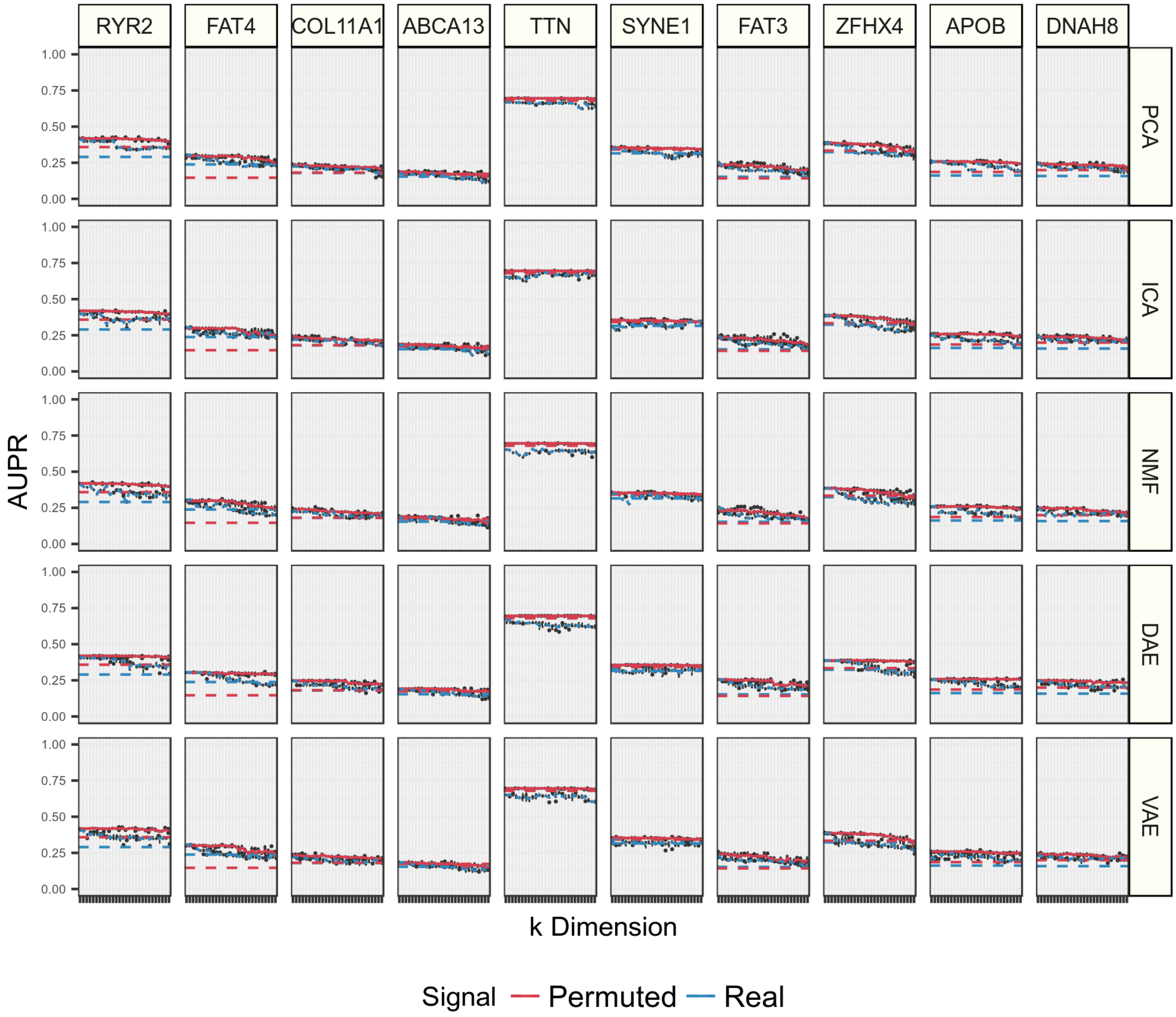
RYR2
FAT4
COL11A1
ABCA13
TTN
SYNE1
FAT3
ZFHX4
APOB
DNAH8
PCA
ICA
NMF
DAE
VAE
AUPR
1.00
0.75
0.50
0.25
0.00
k Dimension
Signal
Permuted
Real

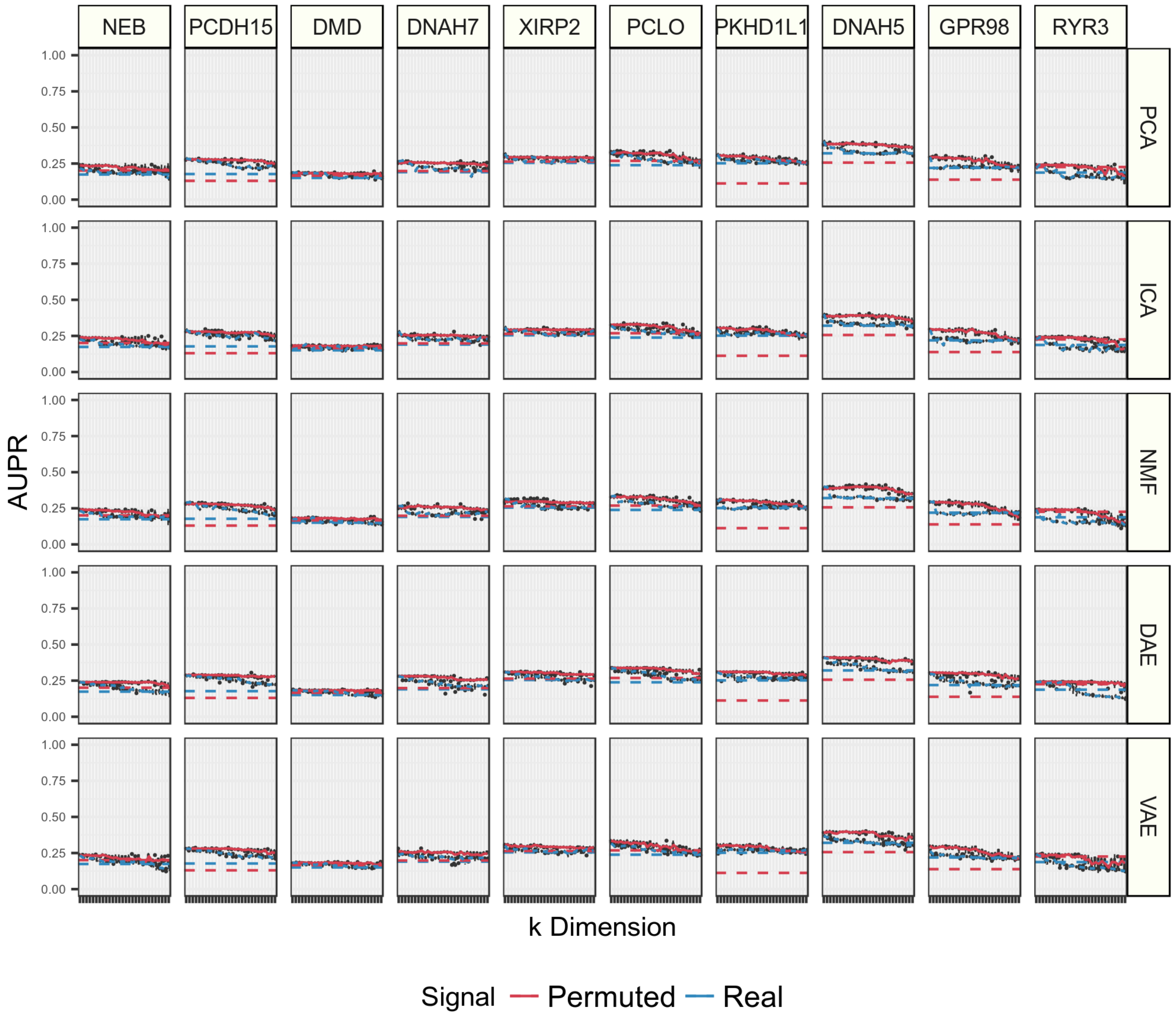

**Figure S13:** *Predicting the top 50 most mutated genes in The Cancer Genome Atlas (TCGA) PanCanAtlas Project with features derived from different compression algorithms across latent dimensionalities.* We independently optimized logistic regression classifiers using compressed features for real and permuted data derived from five compression algorithms across dimensions. All 50 mutations are split across a series of five figures and are presented in order of performance differences between real and permuted data. The area under precision recall (AUPR) for training data in cross validation intervals are shown. The blue lines represent compressed features derived from real training data input into the compression algorithms. The red lines represent compressed permuted data. The dotted lines in red and blue indicate performance with the raw RNAseq features and serve as baselines. All models, including the permuted data models, include covariates for cancer-type and log 10 mutation burden. The distribution of performance is provided in boxplots for all five algorithm initializations. The lines connect the mean of these algorithm initializations.

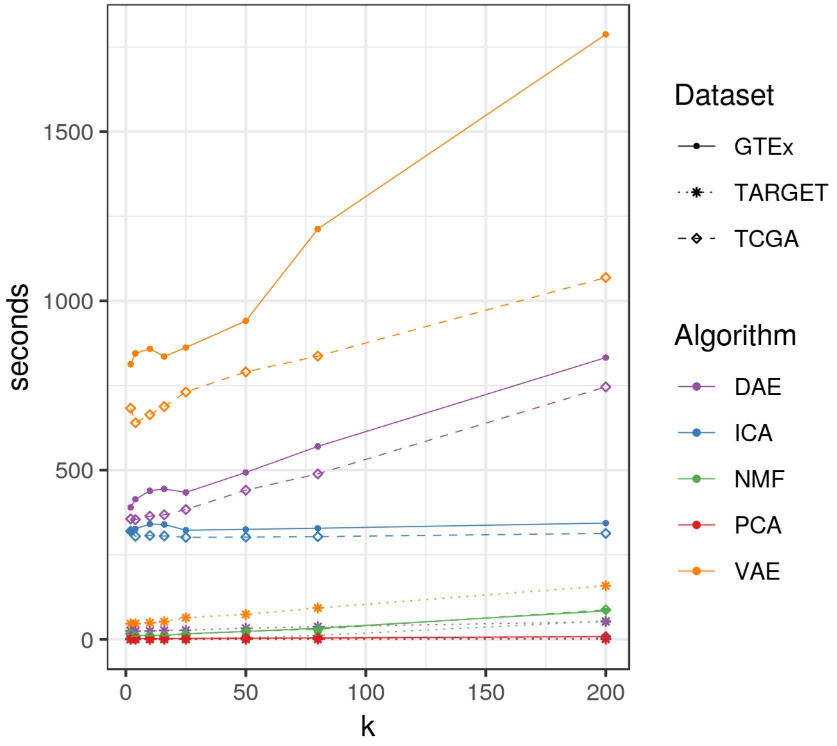

**Figure S14:** *Evaluating execution time of training 5 compression algorithms across latent space dimensionalities.* Comparing time in seconds of principal components analysis (PCA), independent components analysis (ICA), non-negative matrix factorization (NMF), denoising autoencoders (DAE), and variational autoencoders (VAE) in training models on GTEx, TCGA, and TARGET gene expression data. We compared training time across various latent space dimensionalities. Execution time calculated using a machine with an Intel Core i3 dual core processer CPU with 32 GB of DDR4 memory.
