## Supplementary Note for "Sequential compression of gene expression across dimensionalities and methods reveals no single best method or dimensionality"

### *Optimizing Training Hyperparameters in Neural Network Architectures*

We applied BioBombe using five compression algorithms. Two of the five models, variational autoencoders (VAE) and denoising autoencoders (DAE), are based on autoencoder (AE) frameworks and include several hyperparameters that must be tuned to optimize signal reconstruction. Our primary concern in training the AE models was to ameliorate potential performance biases as the bottleneck dimension increased if hyperparameters were kept static. In other words, we sought to isolate performance differences to the effects of changing $k$ dimensionalities. Therefore, we performed a grid search around several hyperparameters for both AE models including 6 representative $k$ dimensionalities.

Training autoencoders, and neural networks in general, requires architectural and hyperparameter decisions to optimize learning important signals in input data. We searched through a grid of various combinations of learning rates, epochs, batch sizes, sparsity, and noise parameters for DAE models, and learning rates, epochs, batch sizes, and kappa values for VAE models. We included 6 representative $k$ dimensionalities in this grid ($k$ = 5, 25, 50, 75, 100, and

125). While many combinations did not converge, we observed relatively consistent performance across all hyperparameter combinations tested for both AE models, especially for the VAE models (**Additional File 2: Figure S2**). We selected the hyperparameter combinations for the top performing models and used these in training downstream models.
